# Deep unsupervised learning methods for the identification and characterization of TCR specificity to Sars-Cov-2

**DOI:** 10.1101/2023.09.05.556326

**Authors:** Yanis Miraoui

**Affiliations:** Stanford University

## Abstract

The T-cell receptor (TCR) is one of the key players in the immune response to the Sars-Cov-2 virus. In this study, we used deep unsupervised learning methods to identify and characterize TCR specificity. Our research focused on developing and applying state-of-the-art modelling techniques, including AutoEncoders, Variational Au-to Encoders and transfer learning with Transformers, to analyze TCR data. Through our experiments and analyses, we have achieved promising results in identifying TCR patterns and understanding TCR specificity for Sars-Cov-2. The insights gained from our research provide valuable tools and knowledge for interpreting the immunological response to the virus, ultimately contributing to the development of effective vaccines and treatments against the viral infection.

## 1 Introduction

The ability to identify T-cell receptors specific to the antigens of a given virus is key to the design of vaccines and therapies. Designing effective vaccines and therapies against a particular virus requires knowledge of which TCRs are specific to the antigens of that virus (Gallagher, 2023). This information can be used to develop vaccines that stimulate the immune system to produce T-cells with those specific TCRs or to develop therapies that use T-cells with those TCRs to target and destroy infected cells. It could, for example, help adoptive T-cell therapies, in which T-cells with specific receptors are infused into cancer patients to target and destroy cancer cells using the specific TCRs. It can also aid in understanding the immune response to the virus and how it evolves over time.

However, identifying the specific TCRs involved in the immune response to a virus can be a complex and daunting task. The world has been experiencing this challenge in particular in the recent past with the emergence of Sars-Cov-2 in 2019:

- Firstly, Sars-Cov-2 is a novel virus, meaning that researchers did not have any pre-existing knowledge about the virus’s antigens or the TCRs involved in the immune response to it. This made it necessary to start almost from scratch to identify the relevant TCRs (Mateus et al., 2020).
- Secondly, TCRs are highly diverse and can vary greatly between individuals. This means that different people can have different TCRs specific to the same virus. Therefore, it is essential to analyze TCRs from a large number of individuals to identify the TCRs that are most commonly associated with the immune response to COVID-19.
- Finally, TCR sequencing is a complex and time-consuming process, which can limit the speed and scale of TCR analysis. However, advances in sequencing technologies and computational methods have now made it possible to analyze TCRs from large numbers of individuals more efficiently (Bravi et al., 2021). Nonetheless, we could take this even further, as modelling the mapping from TCR sequences to binding specificities would allow us to narrow down the number of receptors to test. In other words, we could establish links between TCR sequences and functional properties to help predict binding specificity. This integration would allow promising TCR candidates to be prioritized for experimental validation. This would significantly optimize time and resources.

Despite these challenges, researchers have made some progress in identifying the specific TCRs involved in the immune response to COVID-19. For example, multiple studies have identified TCRs specific to different antigens of the SARS-CoV-2 virus, which can help inform the design of vaccines and treatments. However, this task remains very complex and time-consuming. The exploration of an *automatic* method ^1^ to identify binding TCRs and antigens appears as the leading solution to overcome these barriers.

In the meantime, in the past few years, we have been observing a rapid proliferation of machine learning methods in all domains and particularly in the field of biology. As the *2022 AI Index Report* (Zhang et al., 2022) and *2023 AI Index Report* (Zhang et al., 2023) emphasize, machine learning tools have been democratized exponentially over time, enabling groundbreaking discoveries and accelerating research.

The goal of our study is therefore to find an *automatic* method able to make use of state-of-the-art machine learning tools in order to characterize and identify the antigen specificity of TCRs to Sars-Cov-2.

Multiple research studies have already led to very insightful results in identifying specific immune responses to Sars-Cov-2. In fact, some previous works have been able to relatively well cluster similar TCR sequences based on their antigen specificity using various statistical and machine learning techniques:

- By employing an entirely statistical approach, a custom similarity score based on the frequencies and rates of TCRs can be defined (Altan-Bonnet et al., 2020). For example, some similarity measures are defined using the amino acid sequence of the CDR3 part. This approach can be complemented with foundational clustering methods like k-means or DBSCAN clustering (Cheung, 2022).
- Building a new distance-based TCR repertoire (*tcrdist3*) in order to group TCR based on their metaclonotype (Mayer-Blackwell et al., 2021).
- Using innovative and state-of-the-art deep learning methods in a supervised procedure (Weber et al., 2021) but also an unsupervised approach using AutoEncoders models for example (Sidhom et al., 2021).

Nevertheless, this task is still new and while there is still room to improve the predictive performance of these methods, it also seems essential to analyze the interpretability of these models as it can provide valuable biological insights. The comprehensiveness of these models is crucial because it allows us to understand and relate their predictions to the actual molecular properties and interactions that occur in real life. Furthermore, interpreting the weights, behaviors and internal processes of machine learning methods has become essential to building robust models (Olah et al., 2018).

In our study, we aim to fulfill these precise shortfalls. We intend to compare different unsupervised machine learning methods able to cluster T cell receptors based on their antigen specificity to Sars-Cov-2. Our goal is not only to over-perform current existing methods but also to provide a complete interpretation of the weights and behaviours of these models. We will first analyse the performance of simple models such as the AutoEncoder before building more advanced architectures using a Variational AutoEncoder, Transformers and Transfer Learning.

In our study, we will mainly investigate the antigens linked to the Sars-Cov-2 virus. For this purpose, we will make use of a dataset made freely available here (Nolan et al., 2020). It consists of multiple tabular data files that contain the CDR3, the v-gene and the j-gene sequences of different TCRs (separated by + signs) along with the list of antigens with which they have bound to. This dataset is of a very large size (+300, 000 rows) but for computational as well as vizualisation reasons, we will mostly use some extracts of it. We present a small extract of this dataset in Table 1 to better understand the general structure of the data.

**Table 1:**
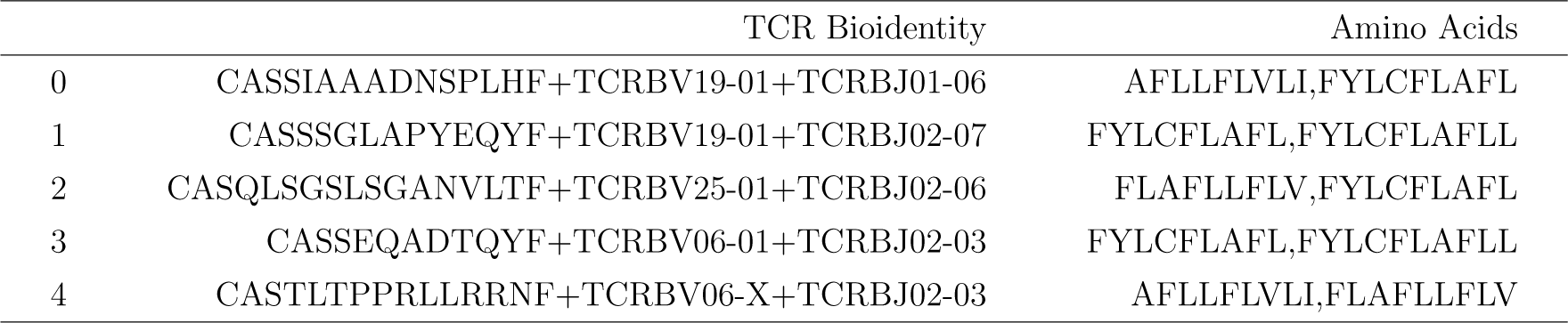
Extract of 5 rows from the dataset; it contains TCR sequences along with the binding list of Sars-Cov-2 antigens.^3^.

Moreover, we will also employ a dataset containing different TCR sequences labeled with their metaclonotype group ^4^ derived from the *tcrdist3* tool (Mayer-Blackwell et al., 2021). Each *.tsv* file in this dataset represents a different metaclonotype group of TCR sequences. We therefore have pre-processed and aggregated it into one file by tagging and numerating each group. This dataset will in particular, help us to evaluate our methods and models and provide valuable tools for the interpretation of our models’ outputs. This dataset is available and can be downloaded from here through the immuneRACE project.

## 2 Results

### 2.1 Comparison of results

In this section we compare the results obtained by applying the different methods and models described in section 4 to our dataset. As mentioned in section ??, the performance of each method is evaluated using three metrics: the Silhouette score, the Calinski-Harabasz index and the Davies-Bouldin index. We compute these metrics as part of our pipeline, following the steps described in the 4.1 section. We then plot our results for the Silhouette score and the Davies-Bouldin index in Figure 1, and provide a comparison table for the Calinski-Harabasz index in Table 3.

**Figure 1:**
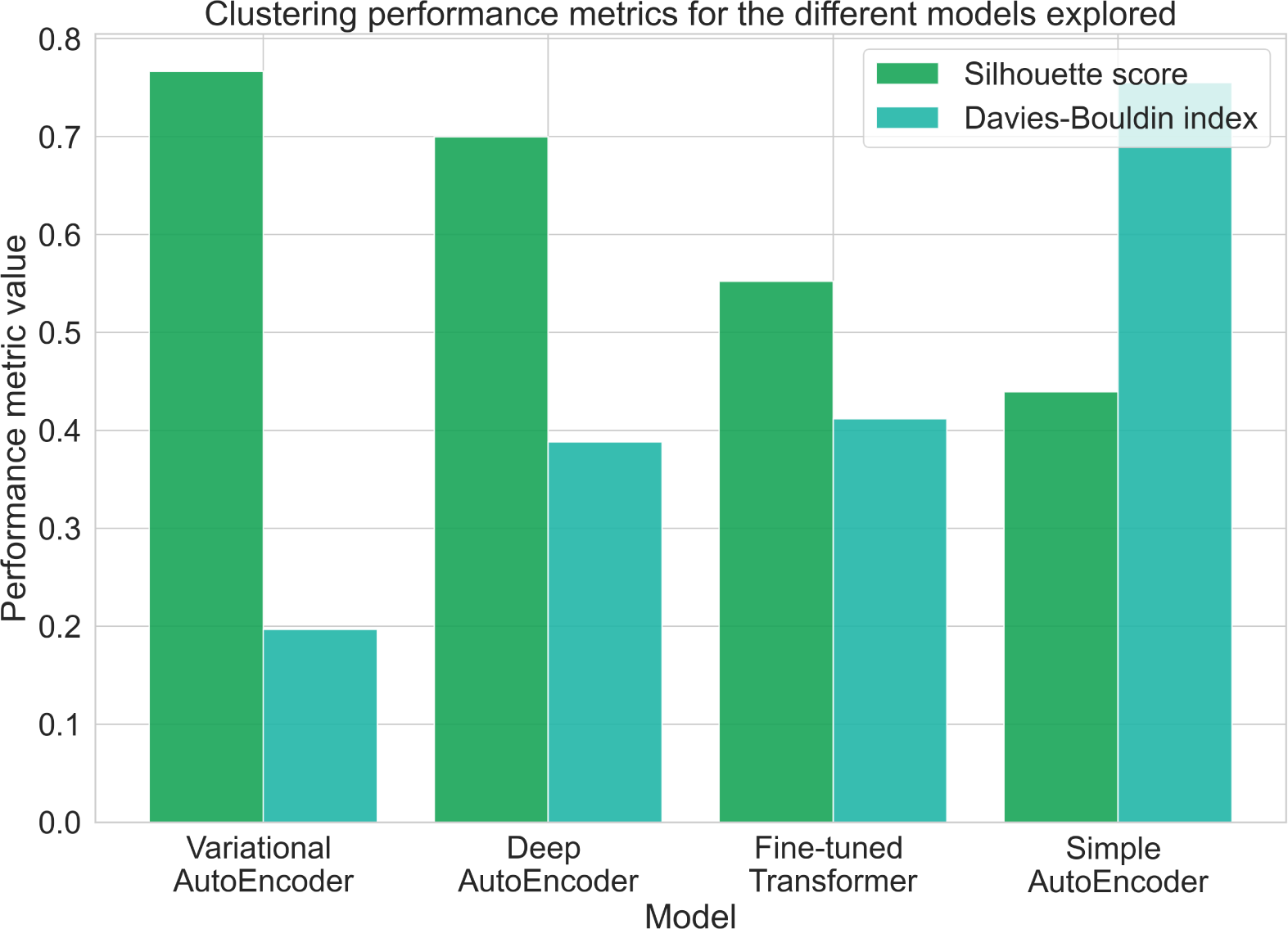
Clustering performance metrics of the models and methods explored in our study.

Looking closely at the results, we can see that the Deep AutoEncoder and the Variational AutoEncoder consistently outperform the Simple AutoEncoder on all three metrics. Both methods achieve higher Silhouette scores, indicating better cluster separation, and lower Davies-Bouldin scores, indicating more compact and well-separated clusters. In addition, the Calinski-Harabasz score is significantly higher for the deep AutoEncoder and the VAE, indicating better cluster quality.

Among the methods, the VAE shows the best overall performance, with the highest Silhouette score and the lowest Davies-Bouldin score. These results suggest that the VAE’s probabilistic encoding contributes to its superior performance in capturing TCR specificity patterns to Sars-Cov-2. Therefore, the VAE represents a promising solution for the effective clustering of TCR sequences.

Furthermore, the TCR-BERT method, which uses transfer learning and transformers, achieves competitive results compared to the other methods. It achieves mod-erate Silhouette and Davies-Bouldin scores and a relatively high Calinski-Harabasz score, indicating its effectiveness in capturing TCR specificity patterns and clustering TCR sequences. It is reasonable to assume that this fine-tuned model could be even more powerful if trained on an even larger dataset.

Overall, the deep AutoEncoder, the Variational AutoEncoder and the fine-tuned TCR-BERT methods show promise in capturing TCR specificity patterns and clustering TCR sequences effectively. However, the VAE stands out as the most successful method, demonstrating its strong potential to accurately and confidently cluster TCR sequences.

### 2.2 Interpretability of our models

Interpretability of deep unsupervised learning models is crucial in the context of identifying and characterizing T-cell receptors specificity to Sars-Cov-2 for several reasons. Here are a few reasons for the importance of model interpretation:

- Interpretable models provide a better understanding of the underlying mechanisms and decision-making processes of the model. This transparency helps to build trust and confidence in the model’s predictions and results. It allows us to validate and verify some of the results of the model, ensuring that the conclusions drawn are reliable. This is even more important for neural networks and deep learning models, which are often seen as *”black-box”* methods.
- Interpretation of deep unsupervised learning models can provide insights into the features or patterns that contribute to TCR specificity. Understanding these patterns can lead to a better understanding of TCR-antigen interactions and inform the design of targeted therapeutics or vaccines.
- Model interpretation helps to understand the generalization and transferability of the learned representations. It allows us to assess whether the model has learned biologically meaningful features that are applicable beyond the training dataset. This knowledge is crucial for deploying the model in real-world scenarios and adapting it to other related problems.

Interpreting deep unsupervised learning models in the context of TCR specificity to Sars-Cov-2 is vital for understanding the underlying mechanisms, gaining insights into TCR-antigen interactions and identifying relevant features and potential biomarkers. That is why, we will carefully study the inner mechanisms of each of our model structures in the following sections.

#### 2.2.1 Interpretability for our AutoEncoder model

In order to evaluate the interpretability of our AutoEncoder model and to gain a deeper insight into its inner workings, we use a separate dataset that we presented earlier in section 1. This dataset contains different TCR sequences grouped into different metaclonotype groups. Metaclonotype groups consist of TCRs that are known to share biochemical similarities. By testing the ability of our model to cluster these biochemically similar TCRs correctly, we aim to assess its ability to capture and represent important features related to TCR structure.

For the purpose of testing, we select three metaclonotype groups that are well balanced. This ensures that we have a representative sample from each group to evaluate the performance of the model. We then use our AutoEncoder model to extract latent representations of the TCR sequences from these selected metaclonotype groups. These embeddings encapsulate the compressed information about the TCRs, capturing their essential features.

To assess the quality of the clustering, we plot the extracted embeddings in a two-dimensional space, as shown in Figure 2. The resulting plot provides a visual representation of the distribution and proximity of the TCRs. If the TCRs belonging to the same metaclonotype group are clustered closely together in this space, it indicates that our AutoEncoder model has successfully learned the relevant structural features solely from the provided inputs (the CDR3 sequences, j-gene and v-gene).

**Figure 2:**
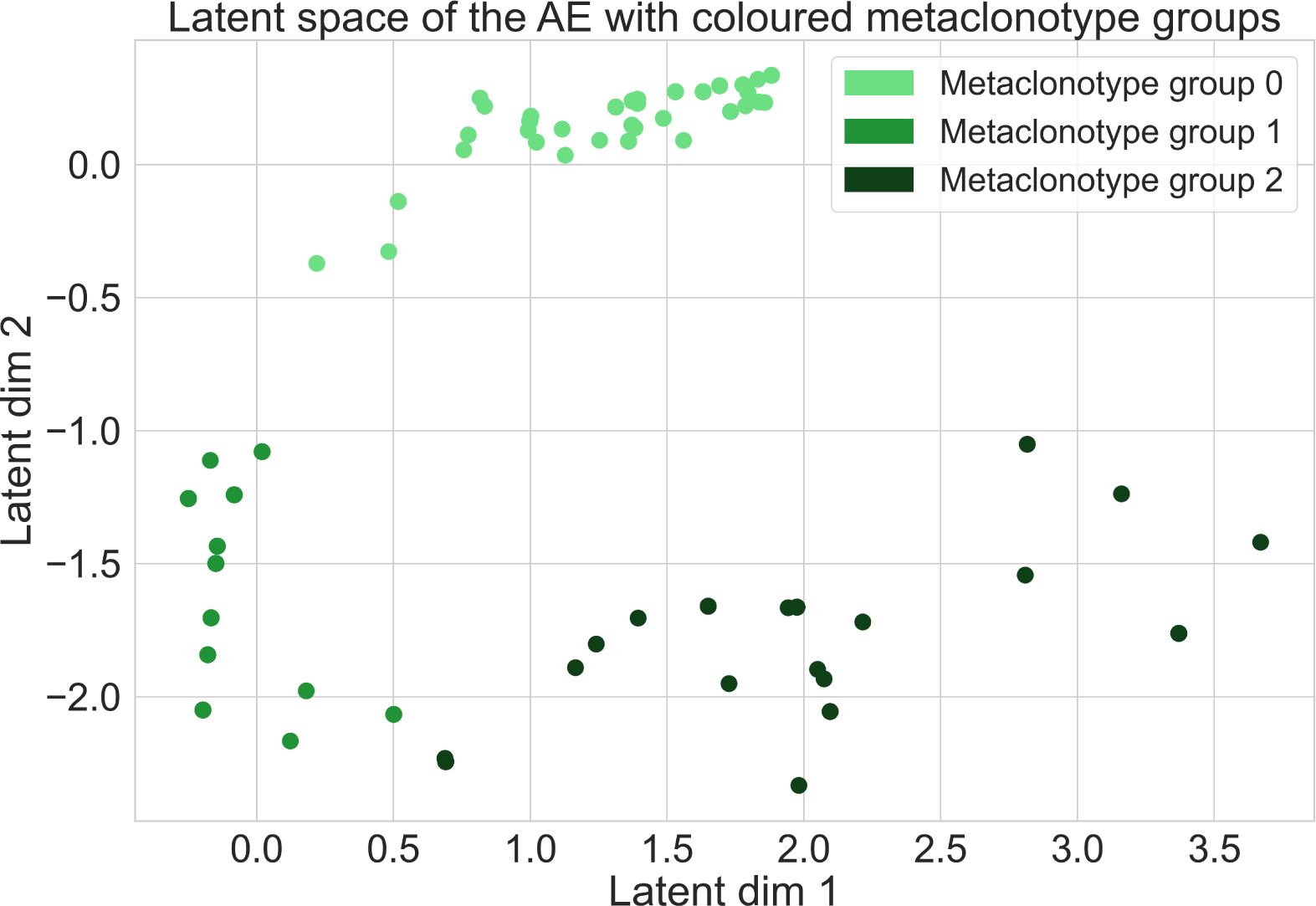
Latent space of the AutoEncoder model with data points coloured with their metaclonotype group.

As we can see in Figure 2, our model is able to cluster biochemically similar TCRs based on these inputs accurately. This means that it has captured meaningful patterns and relationships within the data. This is an important indication of its interpretability, suggesting that it can recognize and represent important structural features of TCRs that are relevant to their specificity for Sars-Cov-2.

#### 2.2.2 Interpretability for our Variational AutoEncoder model

To explore the interpretability of our Variational AutoEncoder model, we focus our analysis on the weight matrices and the variance of the means in latent space. By examining these components, we can gain insights into the patterns captured by the model to differentiate and associate TCRs.

Our first approach is to examine the weights of the encoder component in our VAE model. This examination allows us to understand the relative importance given to each element of the input. As shown in Figure 3, we observe a consistent pattern across specific weight matrices. In particular, we notice that higher weight values are assigned to input positions between 12 and 16. This suggests that the amino acids present in the CDR3 sequence at positions 12 to 16 play an essential role in the differentiation and grouping of TCRs. This finding is consistent with previous scientific research which has shown that the middle part of the CDR3 sequence contains the most variable region and is the main contributor to TCR specificity.

**Figure 3:**
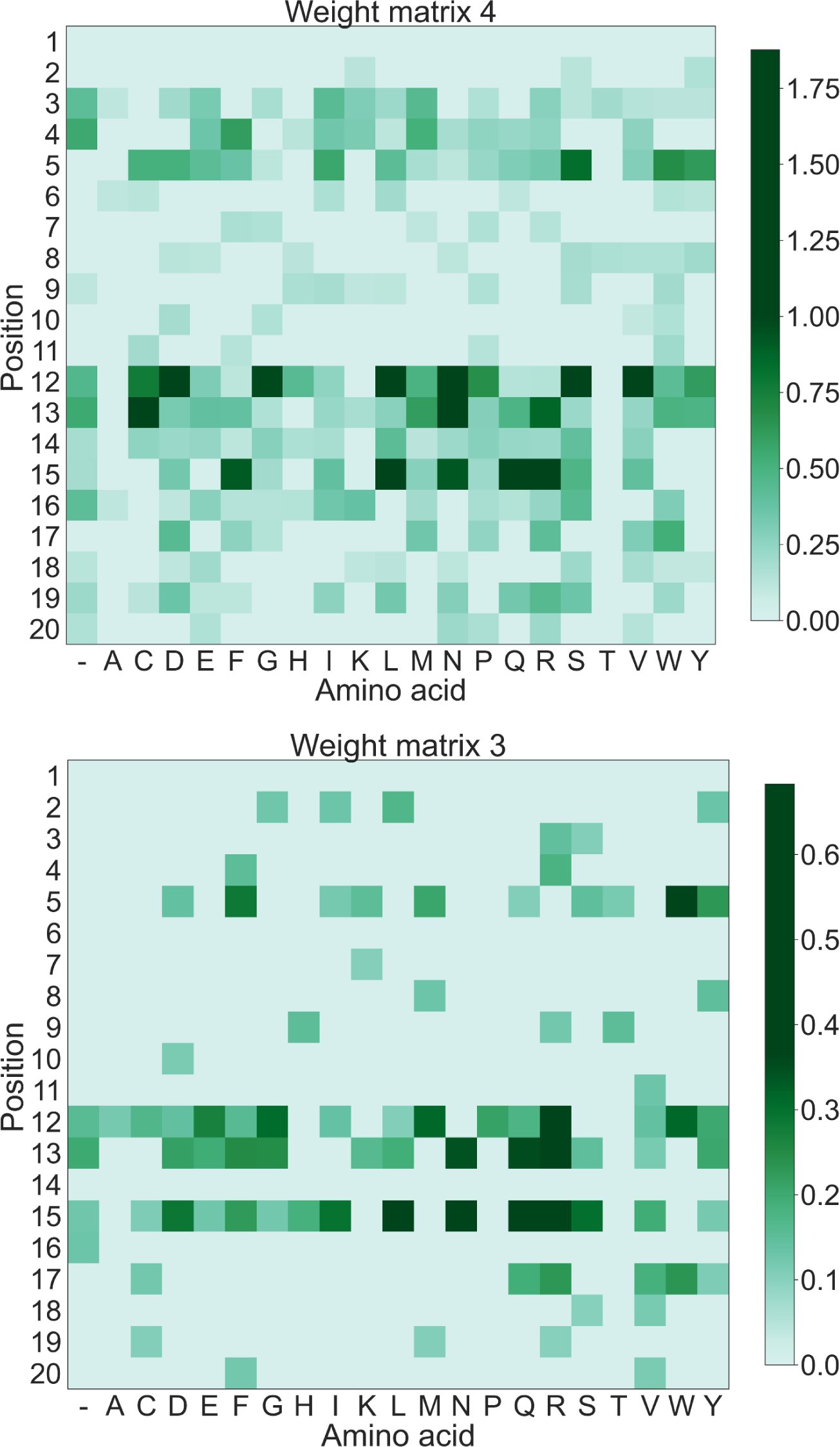
Matrices of the weights 3 and 4 of the VAE model.

By analyzing the weight matrices, we gain valuable insights into the learned representations of the model. The emphasis on specific positions within the CDR3 sequence provides evidence that the VAE has successfully recognized the relevant patterns associated with TCR differentiation and grouping. This understanding improves our interpretation of the model’s behavior and strengthens the link between its internal mechanisms and the underlying biological processes.

Alternatively, if we examine the variance of the means of the distributions in latent space, we observe a similar behavior. To investigate this, we introduce variations in each amino acid at each position and calculate the variance of the means of the resulting embeddings. This approach provides a more intuitive understanding, while giving results that are very close to those obtained by analyzing the weight matrices.

If we plot the variance of the means per position, as shown in Figure 4, we observe a consistent pattern. Once again, we find that the means have a higher variance in the middle of the CDR3 sequence, specifically between positions 10 and 16. This result confirms that our VAE model effectively captures meaningful patterns and relationships present in the data. It serves as an important indication of the interpretability of the model, suggesting that it is capable of recognizing and representing important structural features of TCRs that are relevant to their specificity for Sars-Cov-2.

**Figure 4:**
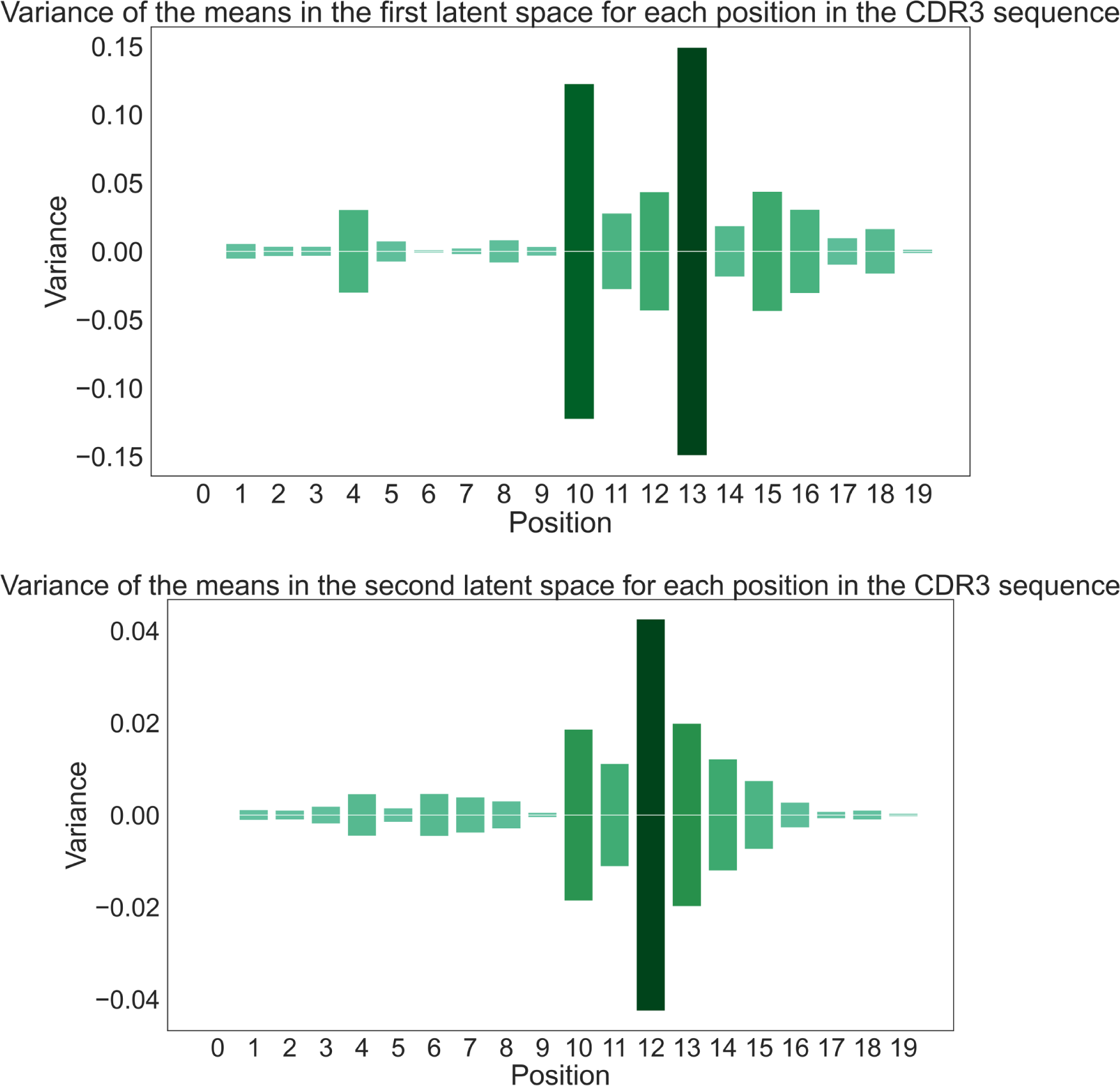
Variance of the means in the latent space at the different positions in the CDR3 sequence.

By thoroughly analyzing both the weight matrices and the variance of means, we gain a comprehensive understanding of how the VAE model interprets and represents TCR data. This insight strengthens our ability to interpret and utilise the results of the model to further our understanding of TCR specificity in the context of Sars-Cov-2.

#### 2.2.3 Interpretability for our Transformer model

In this section, we explore the interpretability of our fine-tuned transformer model, TCR-BERT, by examining its attention heads. By extracting the attention heads from our trained model, we can analyze where the model focuses its *”attention”*. Plotting the attention values for different positions and amino acids, as shown in Figure 5, provides valuable insights and interpretations.

**Figure 5:**
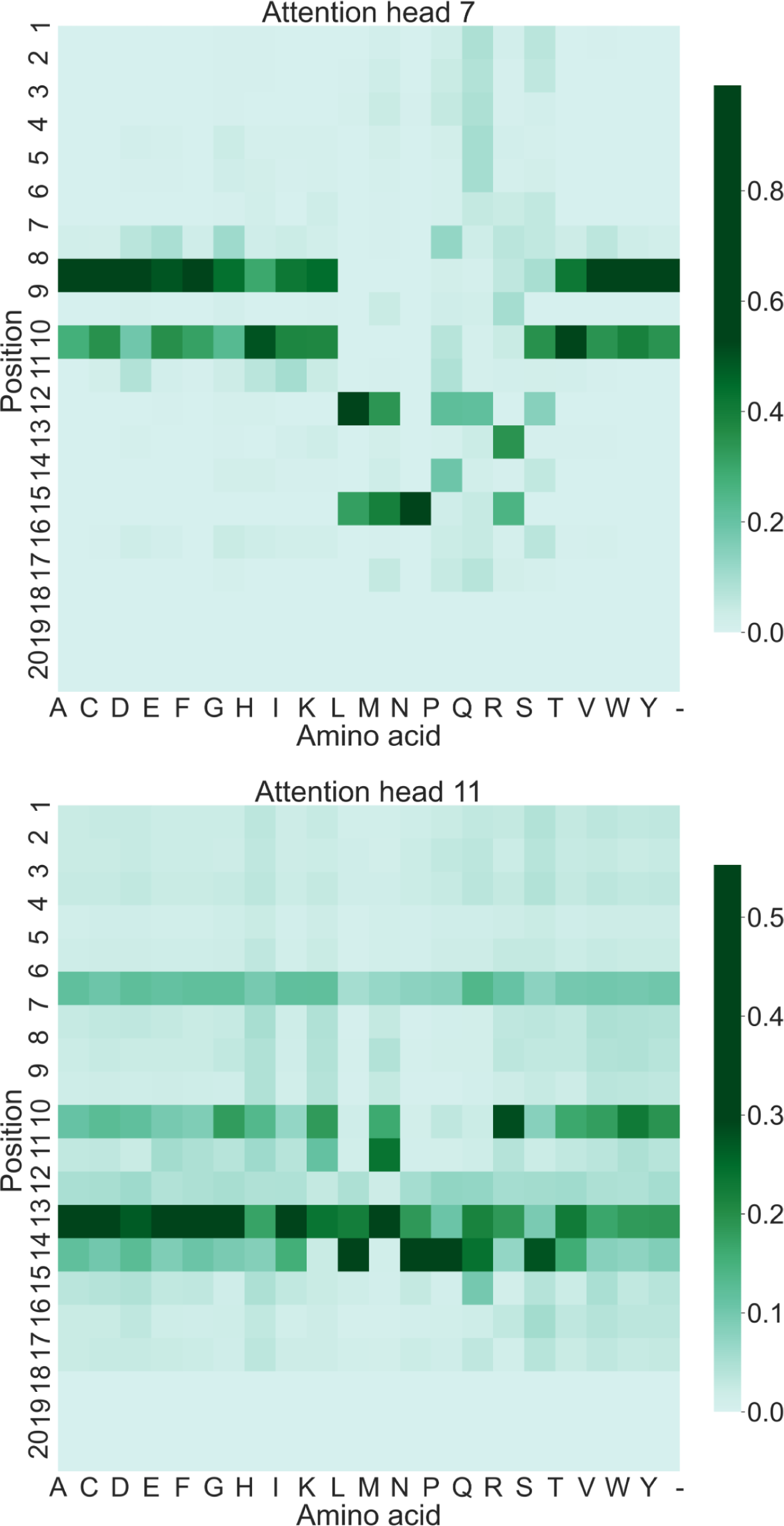
Matrices of the weights for the 7*^th^* and 11*^th^* attention heads of the fine-tuned TCR-BERT model.

Similar to our observations with the VAE model, we find that the attention heads in the transformer model primarily focus on amino acids between positions 8 and 15. This alignment with the VAE model findings is consistent with previous scientific research which, as mentioned above, highlights the middle portion of the CDR3 sequence as the most variable region contributing significantly to TCR specificity.

This result reinforces the notion that both the VAE and Transformer models emphasize the same parts of the input to represent TCRs in a latent dimensional space accurately. The focus on the middle part of the CDR3 sequence, where significant structural variations occur, suggests that both models capture and prioritize By us-ing the attention heads of the transformer model, we gain a deeper understanding of how it processes and attends to different parts of the input. This interpretability allows us to elucidate the mechanisms by which the model distinguishes and represents TCRs. The consistent emphasis on the same regions of the input between the VAE and the transformer model provides further validation of the importance of the middle portion of the CDR3 sequence in characterizing TCRs.

### 2.3 Generating new TCR sequences

In this section we will explore how we can use our previously constructed models, such as the VAE or the fine-tuned Transformer model, to generate new TCR sequences with a desired specificity. This will allow us to explore the vast sequence space and potentially discover novel TCRs with solid affinity and specificity for, for example, Sars-Cov-2 antigens. This can help in the development of targeted therapeutics and vaccines. In addition, the generation of new sequences can help to understand the underlying mechanisms of TCR recognition and specificity by studying the patterns and motifs present in the generated sequences.

First, to generate new TCRs using our fitted VAE model, we can make the assumption that our latent space distribution is continuous as explained earlier in section 4.4.2. We can therefore sample vectors from our latent space distribution and then use our decoder to generate meaningful amino acid sequences. Similarly, for the fine-tuned TCR-BERT, we can use the generative capabilities of the Transformer architecture to generate new TCR sequences. By using the learned representations in latent space, we can sample latent vectors and pass them through the decoder component of the model. This process allows us to reconstruct the corresponding TCR sequences, incorporating the desired specificity and structural features.

Several criteria can be used to evaluate the quality of sequences generated from our VAE and Transformer models. One important aspect is to assess the diversity of sequences generated. A high-quality model should be able to generate a diverse set of TCR sequences representing a wide range of potential specificities. This can be assessed by exploring our latent space and generating sequences based on random points from this space. Figure 6 visually demonstrates the rich diversity of CDR3 sequences present in the latent space as each color corresponds to a different generated CDR3 sequence. This analysis provides evidence of the model’s capability to produce diverse and unique sequences, reinforcing its potential for generating novel TCRs with specific specificities.

**Figure 6:**
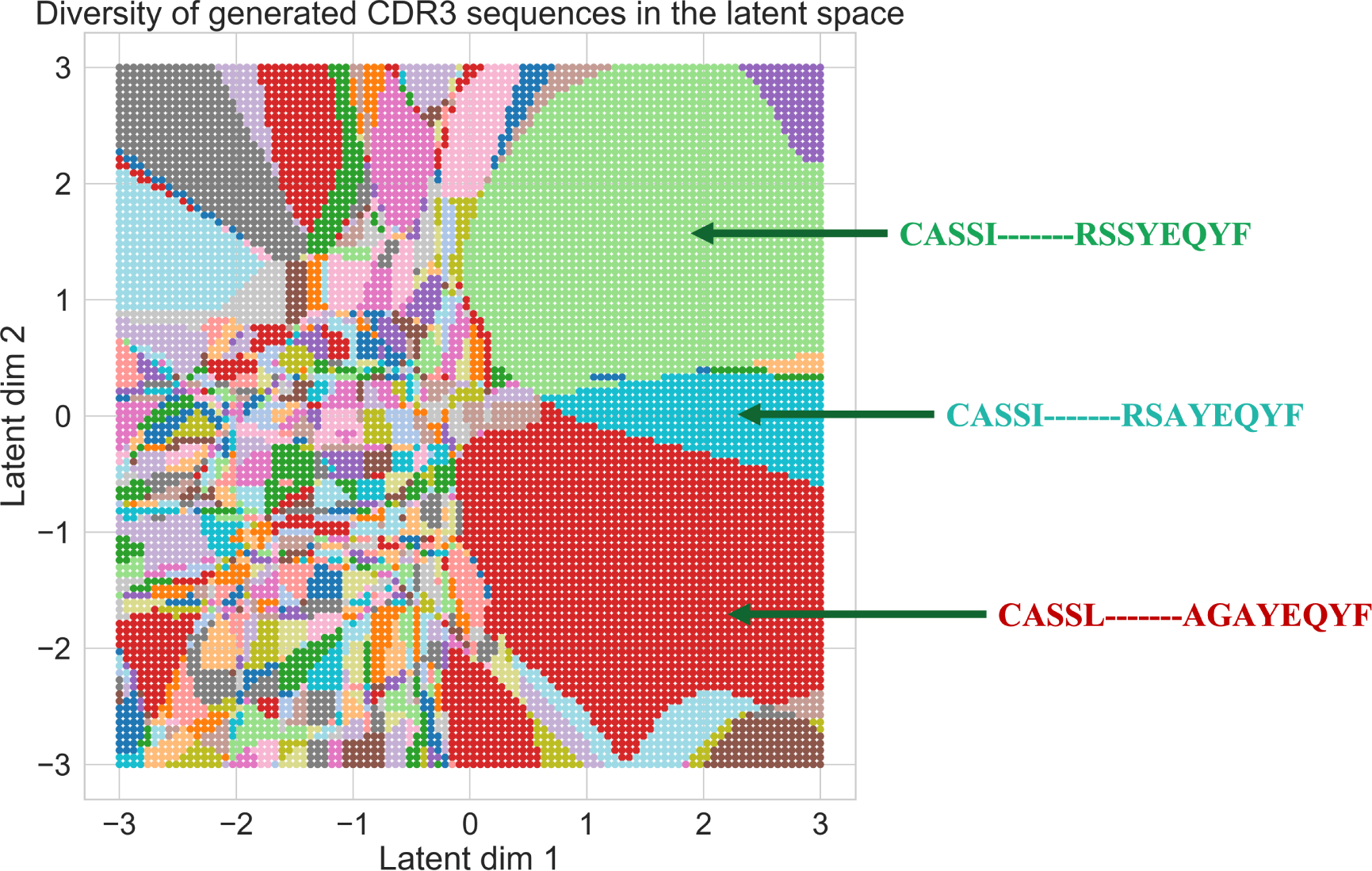
Plot showing how diverse is our latent space in terms of generated CDR3 sequences. Note that each color represents a different CDR3 sequence.

Another critical factor is the structural and biochemical similarity of the generated sequences to known TCRs that show specificity for SARS-CoV-2. The generated sequences should show similar patterns in terms of conserved motifs, amino acid composition and other structural features known to contribute to TCR specificity. Evaluation methods such as motif analysis can be used to assess the similarity be-tween the generated sequences and the desired specific TCRs. The TCRs generated from the Transformer model have previously been evaluated and shown to be similar engineered sequences to those that exist and bind to the same antigen as shown in Figure 7 (Wu et al., 2021).

**Figure 7:**
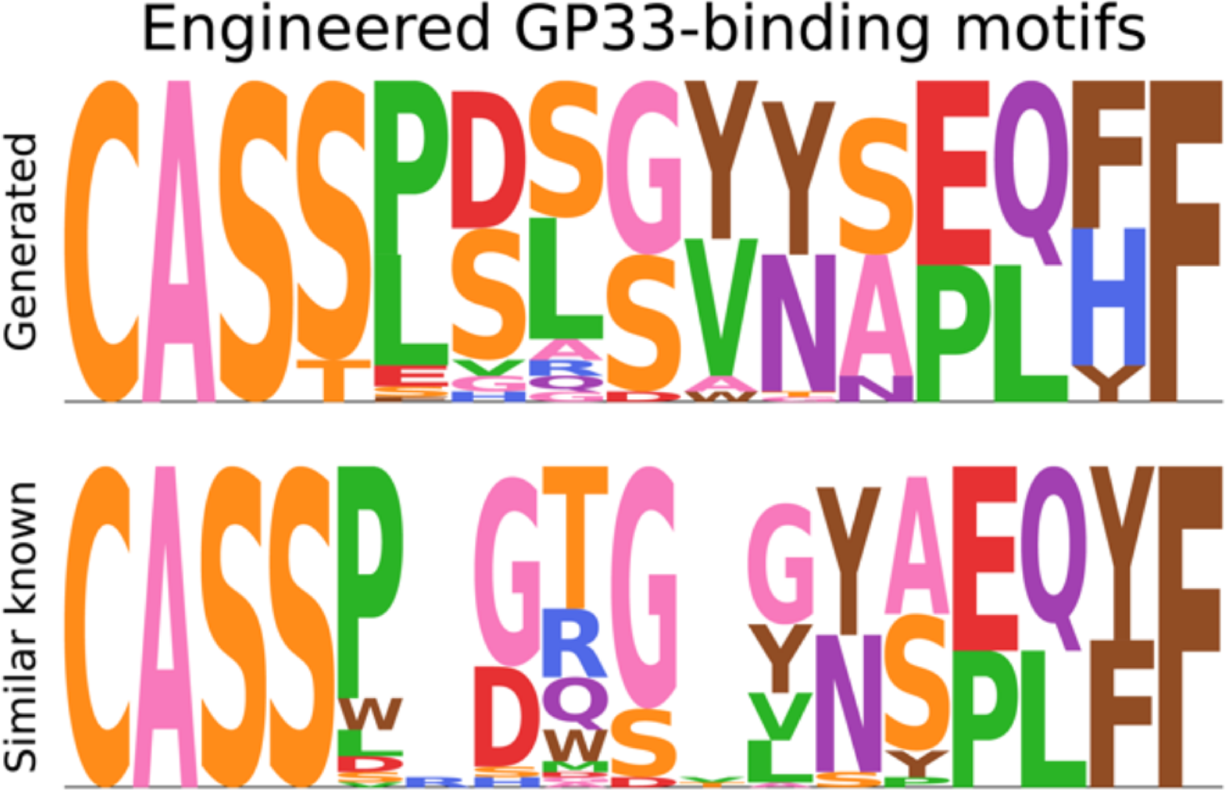
Example of sequences generated using the transformers TCR-BERT along with the motif of similar known sequences that bind to the same antigen GP33 (Wu et al., 2021)

In addition to evaluating the quality of the generated sequences, it is also essential to study the probability of generating these TCRs. The probability distribution associated with the generation of TCR sequences provides insights into the likelihood of obtaining specific sequences. In order to produce these *probabilities of generation*, we leverage a computational tool called **OLGA**^5^ (Sethna et al., 2019). OLGA is developed to compute the generation probability of amino acid and in-frame nucleotide CDR3 sequences from a generative model. Using this tool, our evaluation method is then relatively straightforward:

1. We evaluate the *generation probabilities* of the *”real”* sequences present in our dataset in order to obtain the true distribution of the sequences from our dataset.
2. We then evaluate the *generation probabilities* of the **generated** sequences from our model in order to obtain the probability distribution for the *in-silico* TCR sequences.
3. Finally, we compare the two distributions as shown in Figure 8. If the two distributions are close, it could mean that our generated sequences could have also been part of our initial dataset. This is the case in our study: both distributions seem to be very similar and very close.

**Figure 8:**
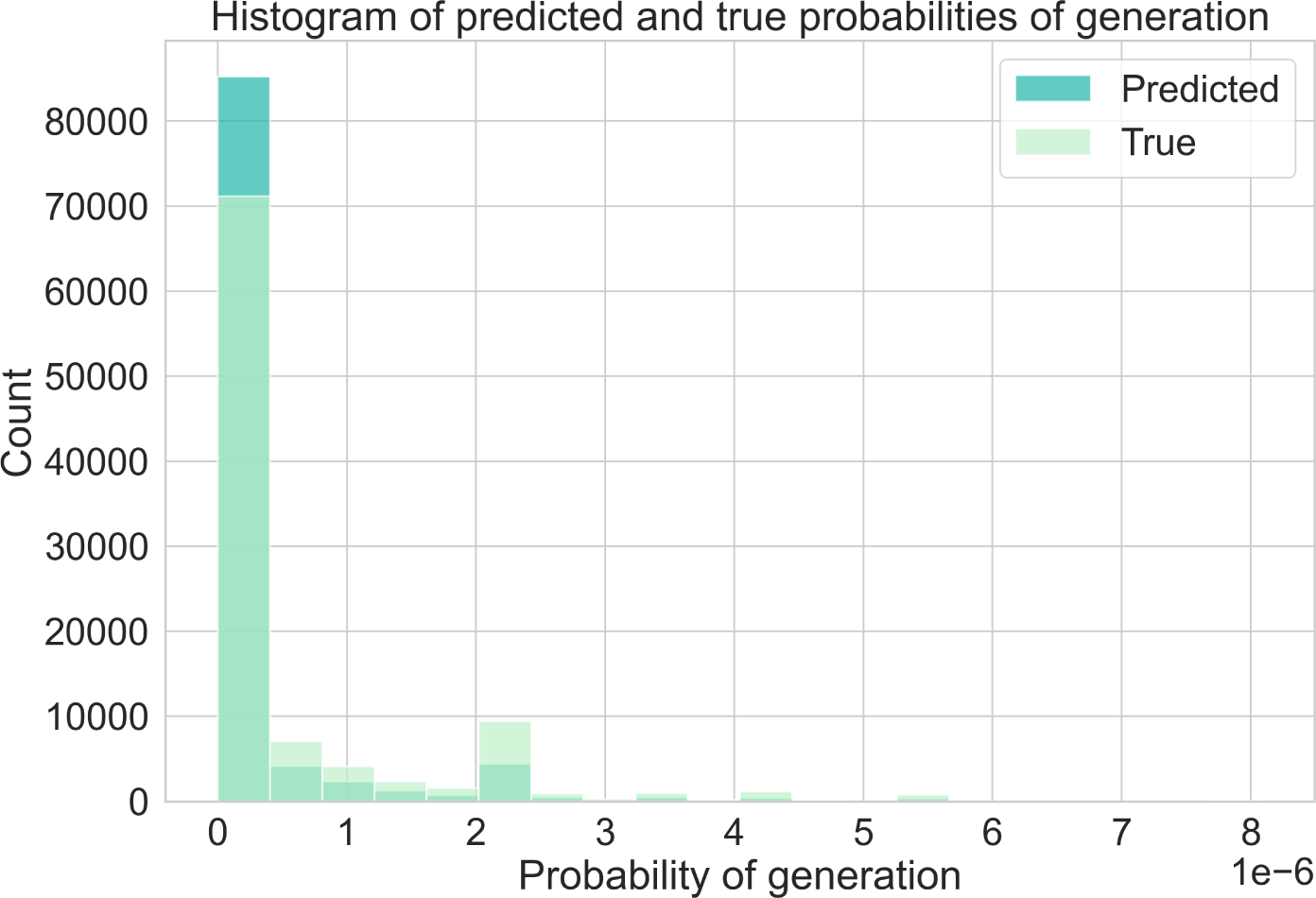
Histogram of the probability distribution for generating the TCR se-quences using the fitted VAE and OLGA.

**Figure 9:**
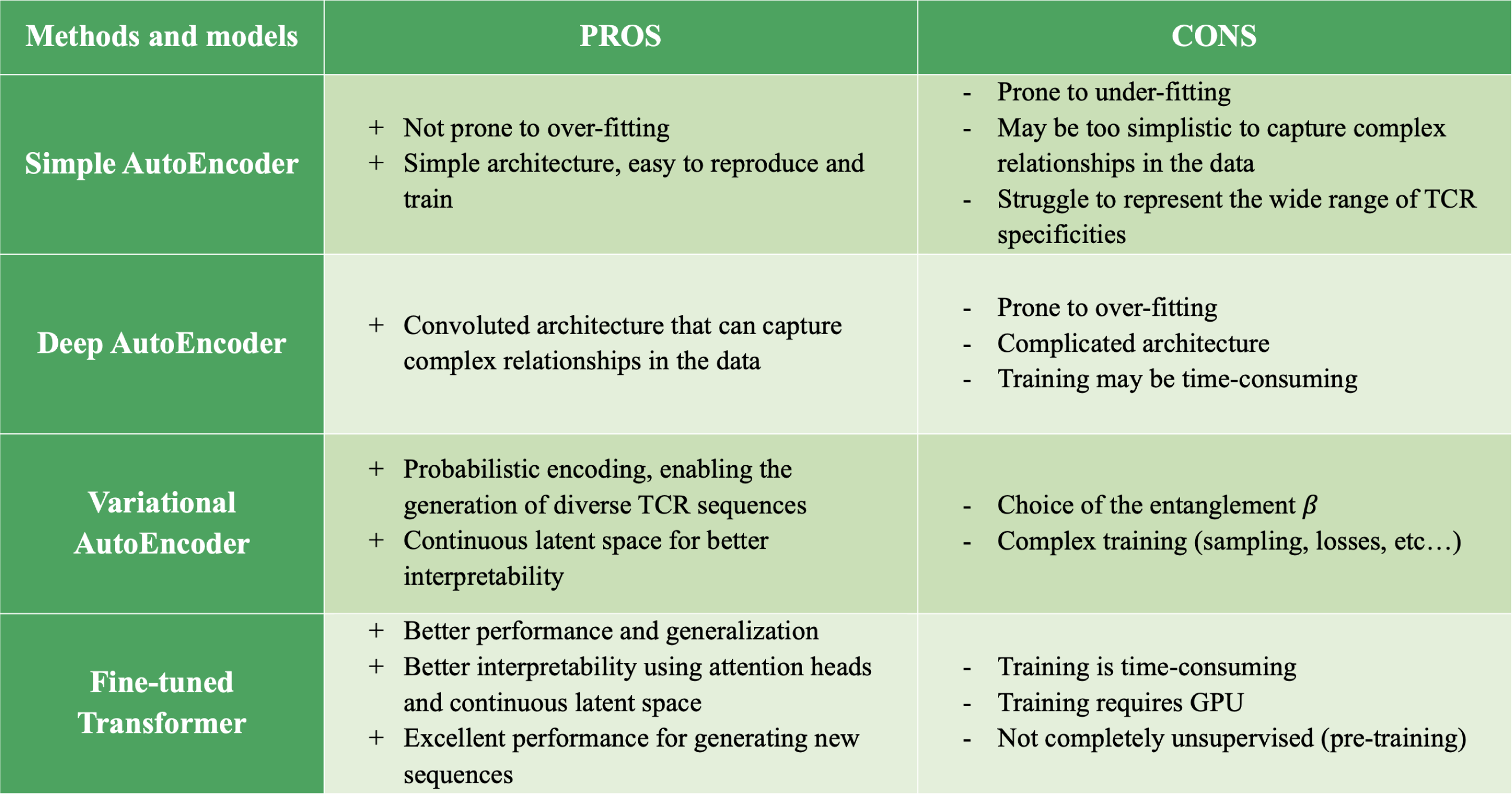
Summary of the pros and cons of the different modelling methods and techniques explored from a data science viewpoint.

It is also important to consider the predicted binding affinity of the generated se-quences to Sars-Cov-2 antigens. Practical experience can be used to estimate the binding affinity and assess whether the generated sequences are likely to have the desired specificity. As this evaluation would require *”hands-on”* biological experiments, we are not able to perform these tests in our study.

## 3 Discussion

In this study, we used deep unsupervised learning methods to identify and characterise T-cell receptor specificity for the Sars-Cov-2 virus. Our research focused on the development and application of state-of-the-art modelling techniques, including AutoEncoders, Variational AutoEncoders and transfer learning with Transformers, to analyse TCR data.

Through our experiments and analyses, we have achieved promising results in identifying TCR patterns and understanding TCR specificity for Sars-Cov-2. Our models have demonstrated their effectiveness in capturing meaningful representations of TCR sequences and clustering them based on their similarities. In addition, the interpretability of our models has provided valuable insights into the features and mechanisms underlying TCR recognition and response to the virus.

Furthermore, our study has highlighted the potential of deep learning methods in TCR analysis for viral infections, such as Sars-Cov-2. The application of transfer learning and Transformers, in particular TCR-BERT, has demonstrated the benefits of applying Natural Language Processing techniques to TCR data, opening up new avenues for research and improving our understanding of immune responses.

The outcomes of our research could have significant implications for vaccine and treatment design against Sars-Cov-2. By identifying TCRs that are specifically involved in the immunological response to the virus, we can gain insights into the key targets for vaccine development and the design of therapeutic interventions. Our re-search has explored the exciting possibility of generating new TCR sequences using the insights gained from our deep unsupervised learning models. By understanding the patterns and characteristics of TCRs involved in the immunological response to Sars-Cov-2, we can utilize generative modeling techniques to create synthetic TCR sequences that mimic the desired specificity and functionality.

In conclusion, our study has demonstrated the effectiveness of deep unsupervised learning methods in characterizing and identifying TCR specificity to Sars-Cov-2. The insights gained from our research provide valuable tools and knowledge for interpreting the immune response to Sars-Cov-2, ultimately contributing to the development of effective vaccines and treatments against the virus.

While our findings are promising, there are still areas for further exploration. The following are some potential next steps to consider:

- As the availability of TCR data continues to increase, the inclusion of more extensive and diverse datasets can improve the generalisability and robustness of our models. Including data from different populations and diseases (e.g. influenza) could provide an even more comprehensive understanding of TCR specificity for Sars-Cov-2 and its variants.
- Further optimization of our deep learning models is essential to improve their performance and interpretability. Exploring different other architectures, hyperparameters and optimization techniques could further improve the accuracy and efficiency of TCR specificity prediction. We could for example try to fit a neural network with a different architecture (RNN, GNN, etc…). Moreover, the development of more advanced explicable AI approaches could provide additional insights into the learned representations and decision-making processes of our models.
- While our deep learning models have shown promise in identifying TCR specificity, experimental validation is critical. Conducting in vitro and in vivo studies to validate the predicted TCR-antigen interactions could provide concrete evidence of their functional relevance. In addition, collaborations with immunologists and clinicians could facilitate the translation of our findings into clinical practice.

## 4 Materials and Methods

### 4.1 One pipeline

Throughout our study, we will make use of **one** similar pipeline. This analogous pipeline will consist of the following:

1. A data pre-processing
2. An encoding of the TCR sequences
3. A modelling and fitting process
4. The leverage of a clustering method

Within this pipeline, all the steps will remain similar except for the *modelling component* which we will modify and adapt to take advantage of different approaches. This will allow us to accurately compare the different modelling techniques used and determine which are better suited to this task.

#### 4.1.1 Data pre-processing

The first step that we have to carry out is the pre-processing of our dataset. This consists of the cleaning and the transformation of our data after having carefully inspected its structure. We mainly focus on the pre-processing of the CDR3 part (the first part of the TCR Bioidentity in our dataset in Tables 1 and 2). In fact, the two other parts consist of the *v-gene* and *j-gene* and these will later be encoded into categories.

**Table 2:**
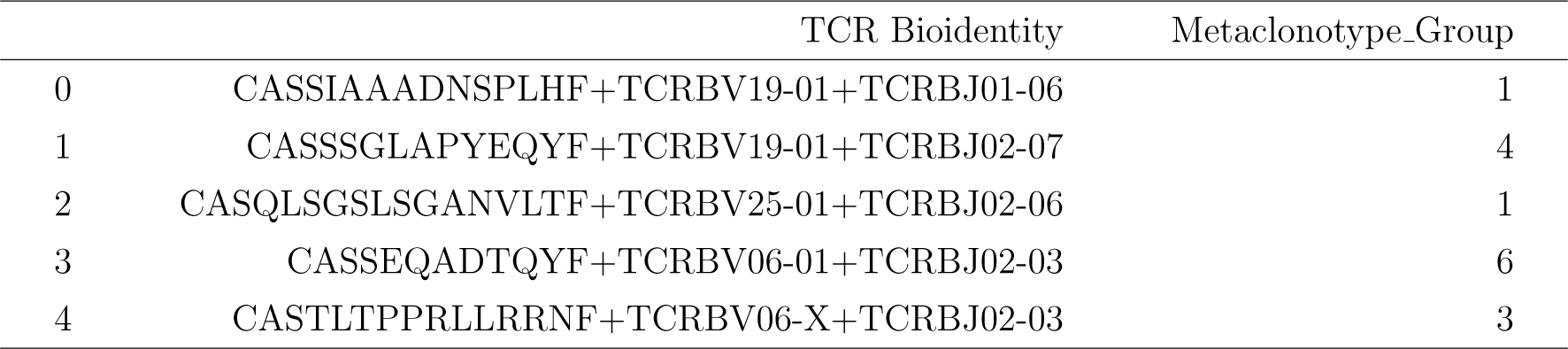
Extract of 5 rows from the dataset; it contains TCR sequences along with their metaclonotype *tag*.

**Table 3:** Calinski-Harabasz index value for each model investigated in our study.

| Model | Calinski-Harabasz index |
| --- | --- |
| Simple AutoEncoder | 1009.71 |
| Deep AutoEncoder | 3769.26 |
| Variational AutoEncoder | 1712.20 |
| Fine-tuned Transformer | 1506.37 |

The first step that we have to conduct is to make sure that there are no errors in the CDR3 sequences of our dataset. In other words, we have to check that each CDR3 sequence that we will be using for our study can be accurately and reliably identified without any ambiguity or inconsistencies. This quality control process is crucial to ensure the integrity of our dataset and the validity of our study’s findings. For this purpose, we employ the computational tools of the *soNNia* package that are designed explicitly for CDR3 sequence analysis (Isacchini et al., 2021). These tools are capable of identifying potential errors, such as inexistent or unrecognized genes, missing values, or other anomalies that could compromise the accuracy of the data.

One of the main obstacles to the exploitation of machine learning methods for the analysis of TCR sequences is the variability in the length of CDR3 sequences. In fact, CDR3 length can vary from a few up to 30 amino acids. We therefore aim to unify this aspect by padding and aligning the sequences according to the procedure presented in a previous study (Bravi et al., 2021). By calculating the length of the CDR3 sequences and the number of amino acids to which the TCR has bound, we are able to have a closer look at the specificities of our dataset. In Figure 10, we can see that the lengths of the CDR3 sequences are symmetrically distributed around 15. By building the boxplot of the lengths of the sequences in terms of the number of antigens the TCR has bound to in Figure 11, we also notice that the lengths of the CDR3 sequences are similar across the different groups of antigens. Only a few *outlier* sequences have a very small (less than 5) or very large (greater than 20) size. Furthermore, we can compute that only **less than 0.4%** have a length greater than 20. Therefore, similarly to the previous study (Bravi et al., 2021), we can choose to align the sequences to all have a fixed length of 20 and drop the small subset of sequences that have larger lengths. The methodology used to align the sequences utilizes Hidden Markov Models (HMMs) and the code is available here. It has also been shown in this previous study that the use of this sequence alignment procedure, with gaps located in the central variable region of the CDR3, does not add any extra bias to our analysis. By using an RBM-LR model that does not require alignment, it has been shown that this alignment routine maintains the known biological structure of CDR3 sequences (Bravi et al., 2021). Therefore, using this alignment process, we obtain CDR3 sequences such as the ones in Table 4.

**Figure 10:**
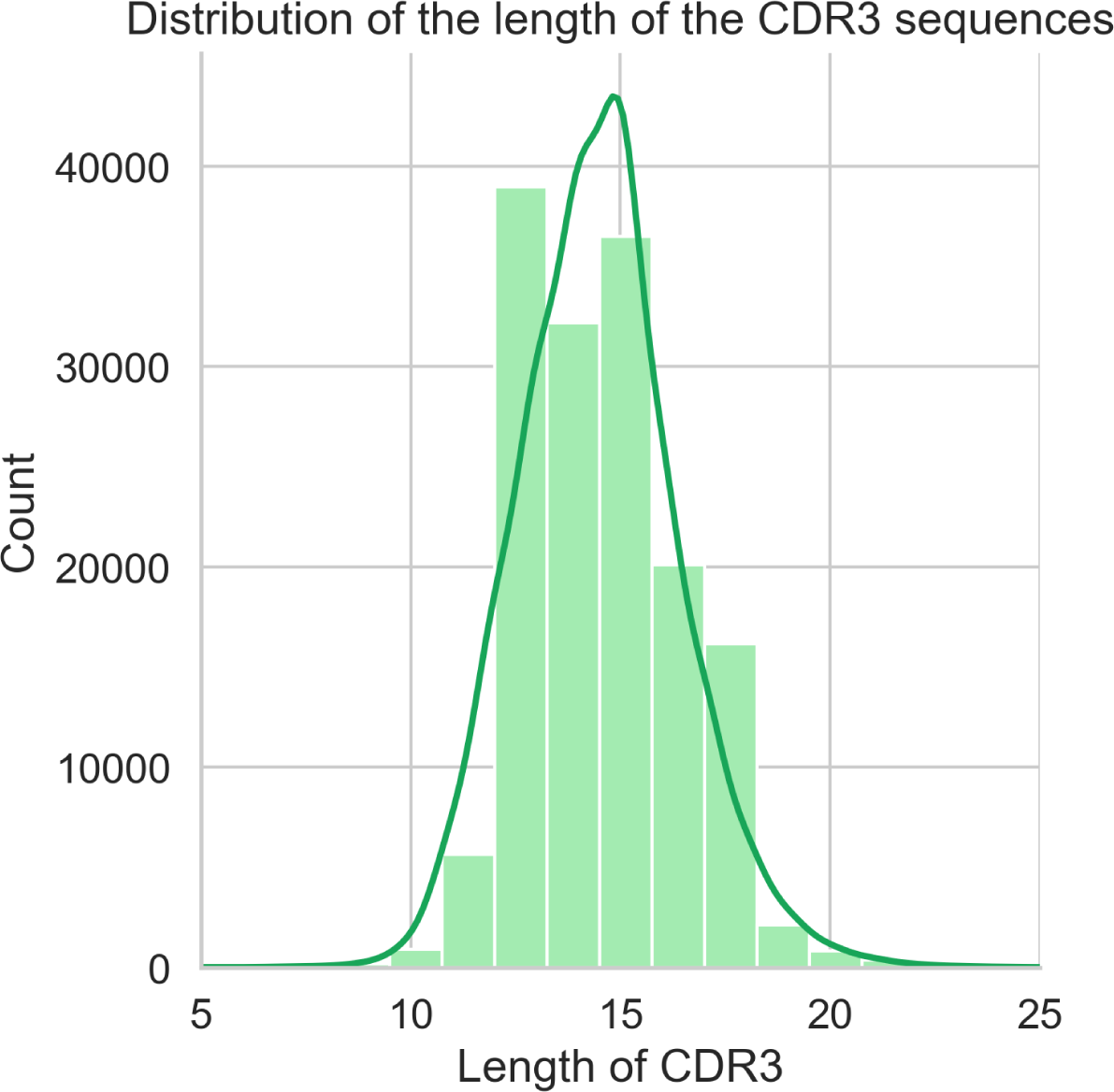
Distribution of the length of the CDR3 sequences.

**Figure 11:**
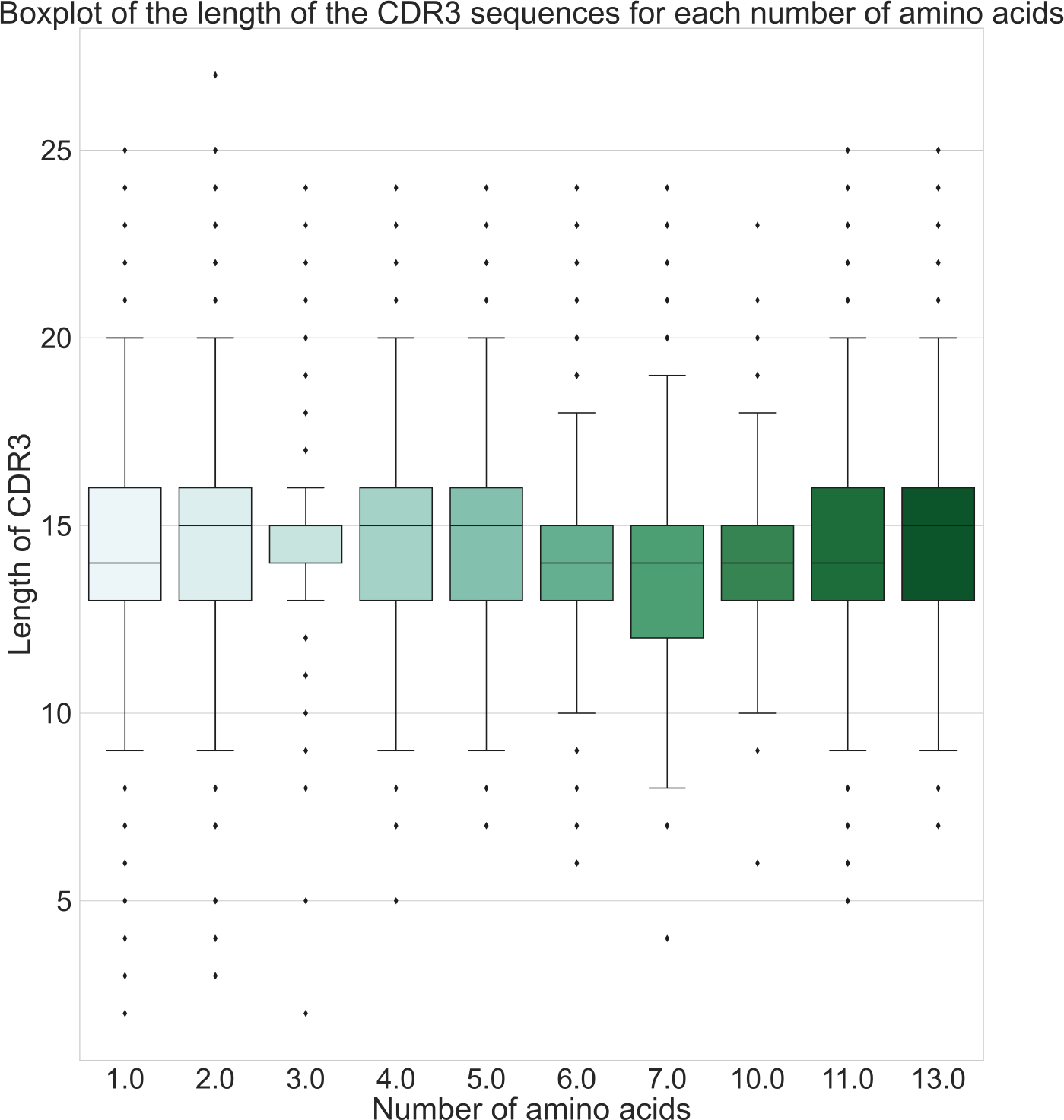
Boxplot of the length of the CDR3 sequences in terms of the number of amino acids the TCR has bound to.

**Table 4:**
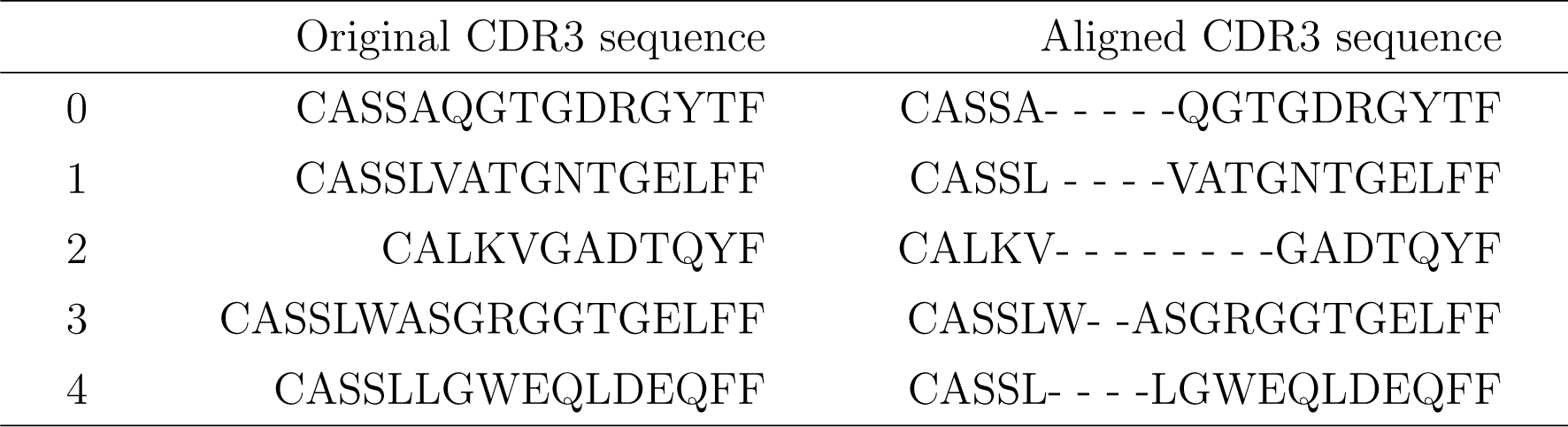
Examples of the result of the CDR3 sequence alignment process.

Furthermore, in order to focus solely on the spatial representation of our TCRs and their antigen specificity clustering, we remove all duplicate rows from the dataset, as the frequency of occurrence is not of interest. Another important aspect that requires pre-processing is the presence of a multi-labeled target for the majority of TCRs. This target is provided as a list of antigens to which the TCRs have bound. However, for our analysis, we only consider the first element of this list as the designated label. By doing this, we aim to evaluate the spatial representations of the TCRs and their clustering based on their antigen specificity to Sars-Cov-2.

#### 4.1.2 Encoding

The next essential step in our pipeline involves the encoding of our CDR3 sequences as well as the *v-genes* and *j-genes* by converting their characters into numerical representations. By converting the sequences and genes into numerical values, we enable the application of various mathematical and statistical techniques for further analysis and modeling of the data.

To encode our CDR3 sequences, we construct an alphabet of all the possible characters that can be present in that string. This alphabet of 21 characters is employed to convert the sequences into one-hot encoded matrices of size 20×21. The alphabet is of size 21 as it is composed of 20 amino acids and the separator **”-”** used during the alignment procedure.

On the other hand, our goal is to make the most of the data available. Given that we have access to both the *v-gene* and *j-gene* sequences, we choose to exploit this additional information as we expect it to improve the performance of our models significantly. To achieve this, we use a one-hot categorical encoder to transform these genes from their character representation into a one-hot categorical format. By using this encoding technique, we convert the *v-genes* and *v-genes* into binary vectors where each gene is represented by a unique category. This transformation allows our models to effectively capture and utilize the distinct features of these genes, potentially improving the overall performance and reliability of our clustering.

Finally, we combine our encoded CDR3 sequences with our two categorically encoded vectors representing the *v-gene* and *j-gene*. As we can see in Figure 12, our input data for our models is therefore a list of 3 matrices. Note that in the following sections, we will refer *input of size 20* to a vector of a sequence that has been encoded into a matrix of size 20 *×* 21.

**Figure 12:**
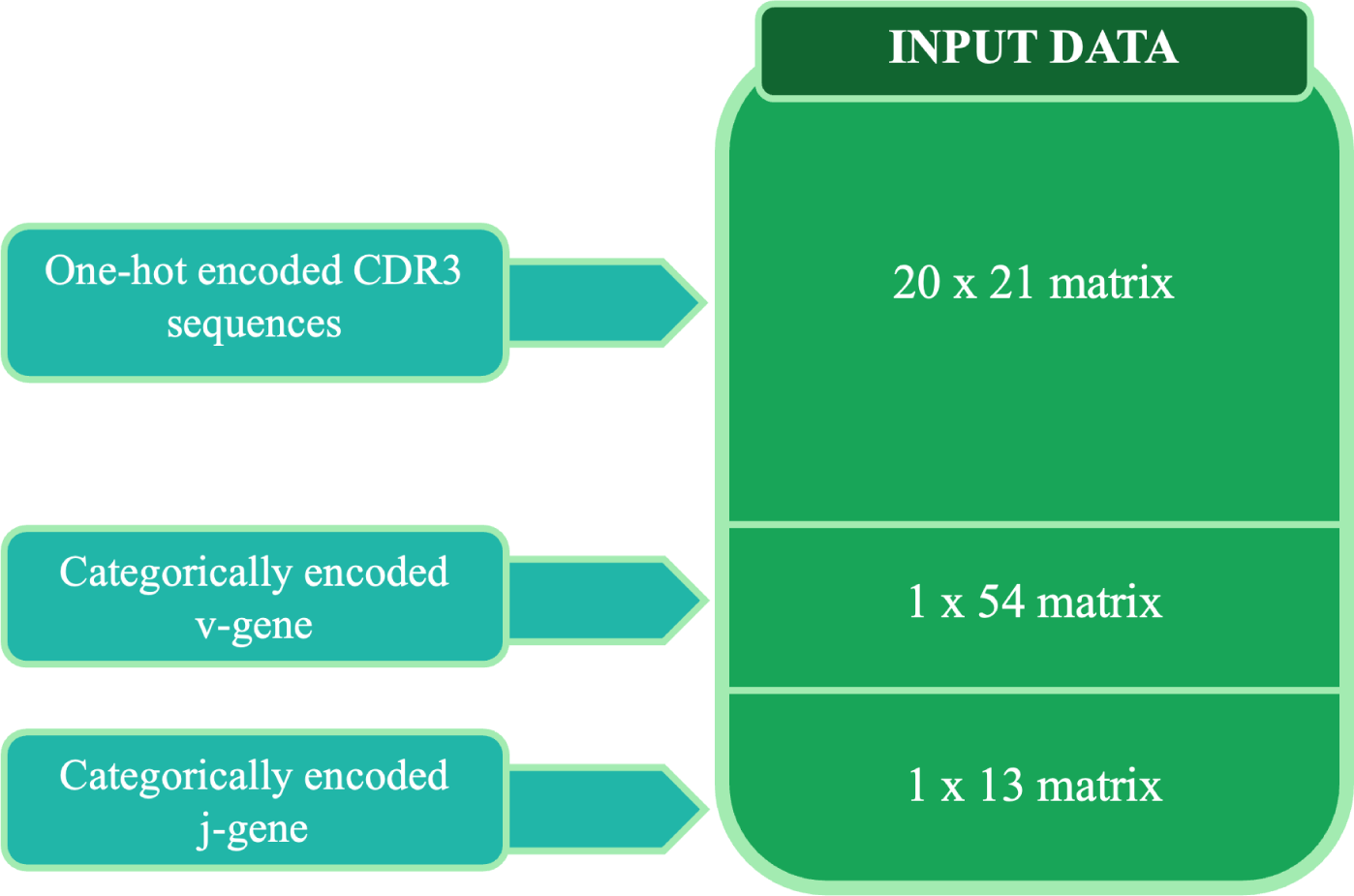
Diagram of the different components of the input data.

### 4.2 A simple AutoEncoder

The first model that we build, fit and evaluate is a simple AutoEncoder model. This seems like a natural choice as the AutoEncoder model is a type of artificial neural network that is commonly used for unsupervised learning tasks. It belongs to the family of feed-forward neural networks and is designed to encode and decode input data with the objective of reconstructing the original input as accurately as possible. Therefore, we will first apply this model to our input data previously defined in Figure 12.

The architecture of an AutoEncoder consists of two main components:

- an encoder *g_ϕ_*
- a decoder *f_θ_*

The encoder network takes the input data and maps it to a lower-dimensional representation, also known as a latent space. This mapping is achieved through a series of hidden layers that reduce the dimensionality of the input. The final layer of the encoder typically produces the compressed representation or code. For the simple AutoEncoder model, we will choose to have one input layer of size 20, one hidden layer of size 40 and a latent space of size 2 as it could be schematize in Figure 14.

**Figure 13:**
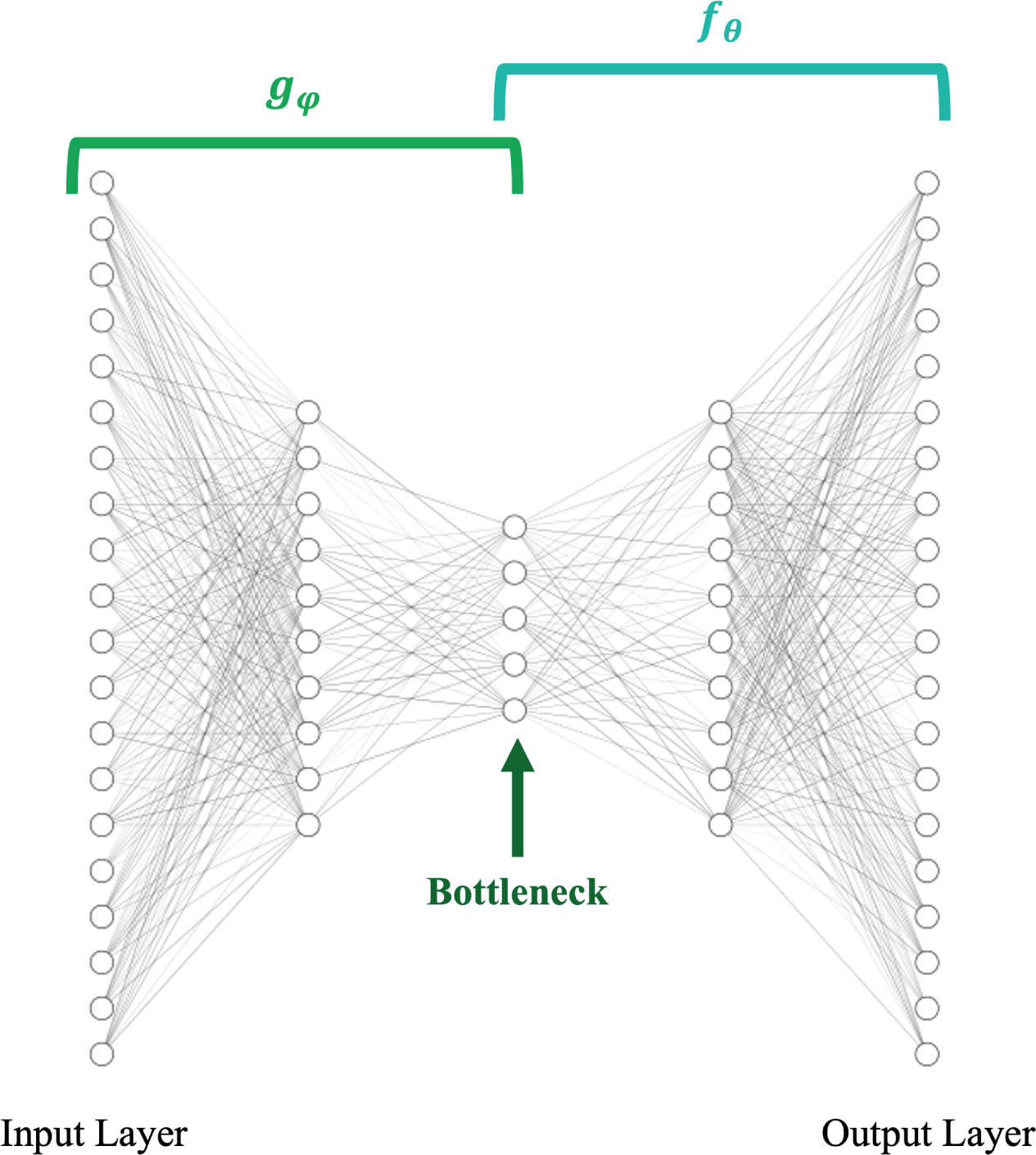
Diagram of a typical AutoEncoder architecture.

**Figure 14:**
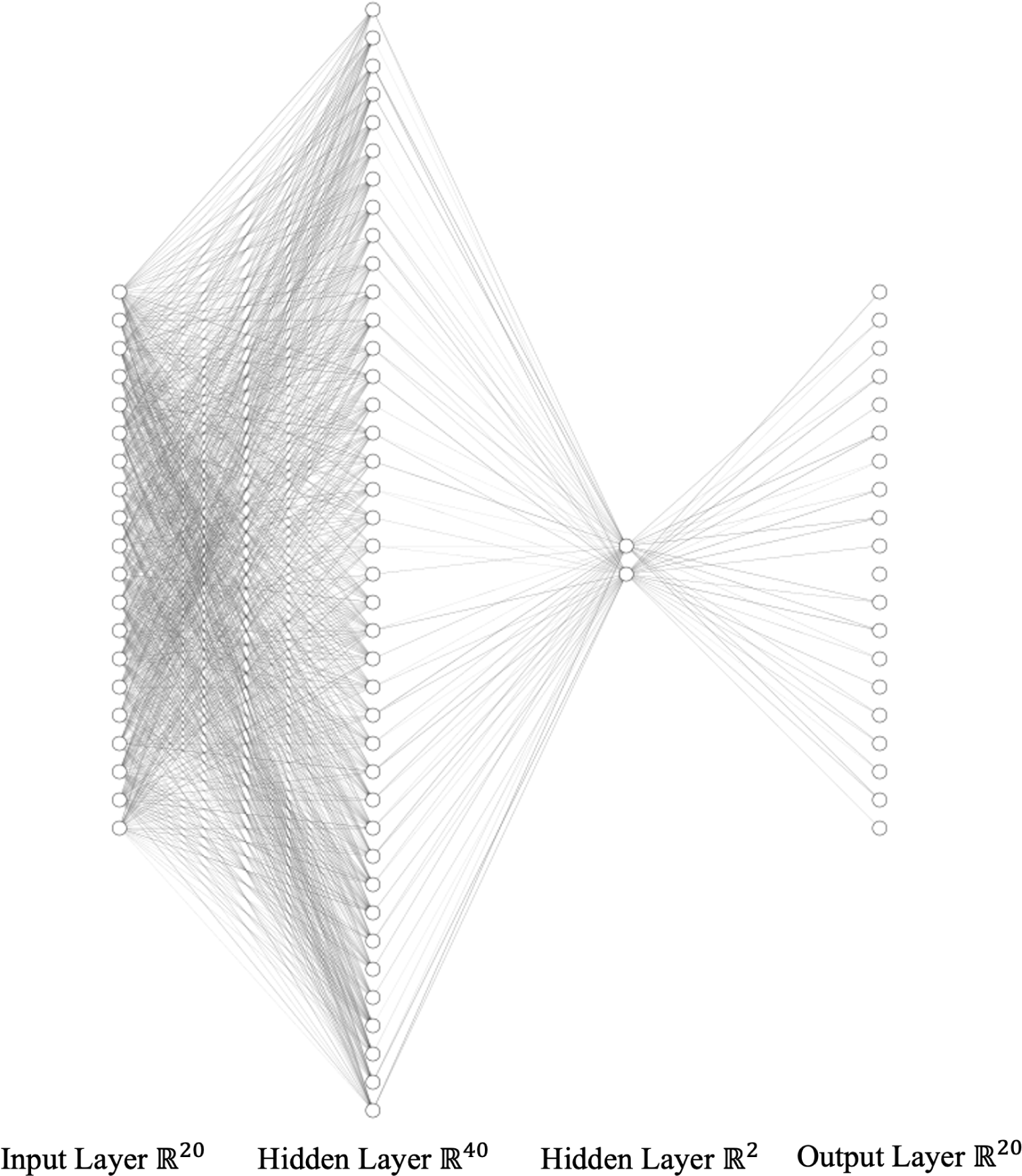
Model architecture of the simple AutoEncoder.

The decoder network takes the encoded representation produced by the encoder and attempts to reconstruct the original input from it. Each layer has an increasing size until the output matches the dimensions of the input. The reconstruction is performed by applying an activation function to the output layer, which generates the output values.

Each layer of our simple AutoEncoder uses **R**ectified **L**inear **U**nit (ReLU) activation functions (Figure 15):

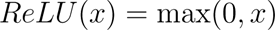

**Figure 15:**
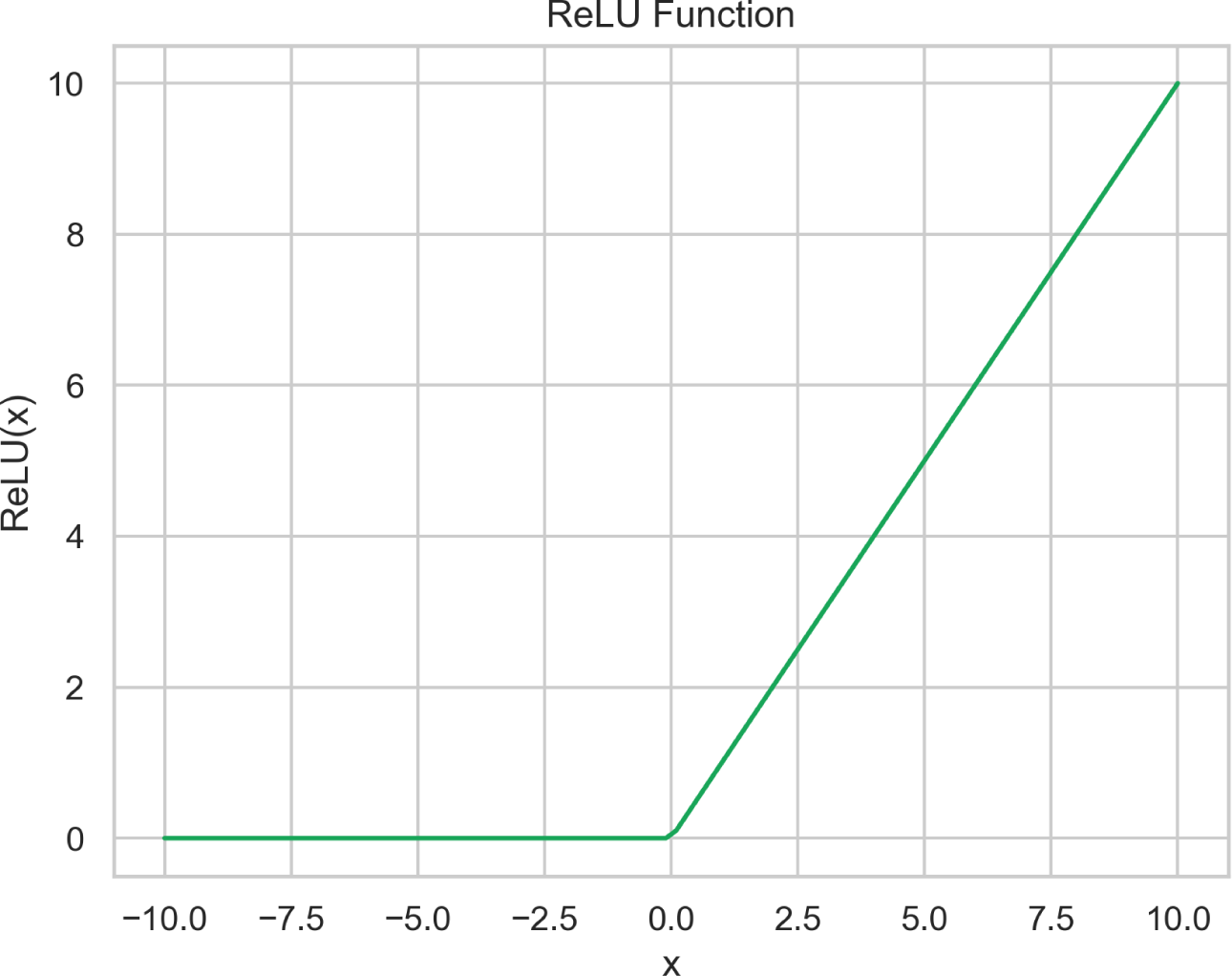
Plot of the ReLU function.

The ReLU activation function is a popular choice for neural networks because of its simplicity and ability to handle non-linearities effectively. It allows the model to capture complex patterns and relationships in the data. In addition, ReLU activations help create sparse representations because they set negative values to zero, effectively suppressing irrelevant or noisy information.

One of the key features of the AutoEncoder is its ability to learn valuable representations or features from the data without explicit supervision. By compressing the input into a lower dimensional space, the AutoEncoder automatically captures the important patterns and structures in the data.

Furthermore, AutoEncoders can also be used for generative purposes, where they are trained to generate new data similar to the input data. By sampling points from the latent space and passing them through the decoder, the AutoEncoder can produce new samples that have similar characteristics to the training data.

### 4.3 An optimized deep AutoEncoder

The previous AutoEncoder model discussed earlier is characterized by its simplicity and smaller architecture size. These uncomplicated models often have significant advantages, such as a reduced risk of over-fitting to the data. However, there is a trade-off as our model may be too simple and may not capture all the information contained in the data due to its limited structure. To address this concern, we decided to construct a larger AutoEncoder model with an optimized architecture and hyperparameters. Although this deeper AutoEncoder model is expected to show some degree of over-fitting to the training data, we may find that its performance exceeds that of the previous model.

As mentioned above, for this iteration, we develop an extended deep AutoEn-coder model with a greater number of layers and larger layer sizes. Another notable difference is the inclusion of additional inputs, namely the v-gene and j-gene of a TCR, in addition to its CDR3 sequence. We expect this inclusion of additional information to have a significant impact on both the complexity and performance of our model, as the v and j genes contain crucial information regarding the specificity of a TCR.

In this iteration, our model is constructed with a dense layer of 87 units, followed by two substantial layers of 75 units each. In addition, the bottleneck of our auto-encoder model has been significantly expanded and now consists of 20 units. Figure 16 illustrates the architecture, where the decoder section consists of a layer of 40 units followed by an output layer of size 20.

**Figure 16:**
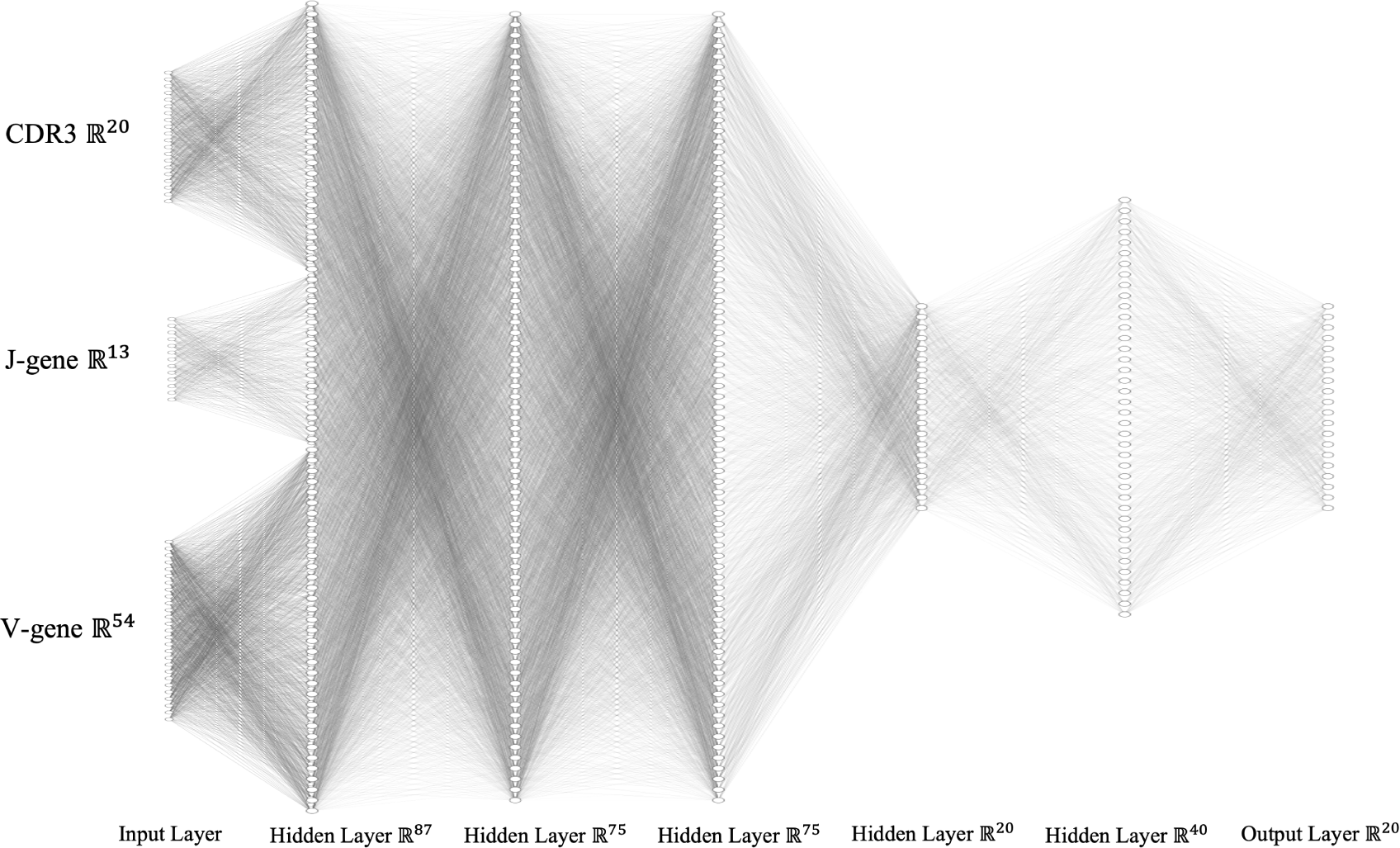
Model architecture of the optimized deep AutoEncoder.

### 4.4 A Variational AutoEncoder

The **V**ariational **A**uto**E**ncoder (VAE) is a type of generative model that combines concepts from traditional AutoEncoders and variational inference. It is also specifically designed to generate new samples that resemble the input data distribution.

Similarly as in the classic AutoEncoder, in a VAE, the encoder network takes an input data point and maps it to a latent space representation. This time, however, this lower dimensional bottleneck is characterized by a mean and a variance. In essence, the encoder learns to capture the salient features of the input data and represents them in the form of a probability distribution in the latent space (Kingma and Welling, 2022).

One distinctive feature of the Variational Autoencoder is its emphasis on imposing a normal distribution structure on the latent space. This is achieved by incorporating the **K**ullback-**L**eibler (KL) divergence term into the VAE’s objective function. By including this term, the VAE encourages the learned latent space to follow a desired prior distribution, typically a standard Gaussian. This constraint ensures that the latent representations have a smooth and continuous nature, allowing for meaningful interpolations and a more interpretable latent space. The VAE’s focus on enforcing normal distributions in the latent space enables it to capture the underlying statistical properties of the data, facilitating the generation of diverse and realistic samples. This also facilitates the effective clustering of the data points by promoting normal distributions in the latent space. This clustering property enables the VAE to capture underlying patterns and structure in the data, further enhancing the model’s ability to generate coherent and meaningful samples based on different modes of the data distribution.

In general, the VAE model offers several advantages: it allows for unsupervised learning of complex distributions, facilitates meaningful latent space interpolations, and supports the generation of novel samples. This is particularly relevant for our study as this model architecture could greatly improve our embedding and clustering performance.

In general, a Variational AutoEncoder (VAE) has a similar structure to the one of a traditional AutoEncoder. The architecture of a VAE consists of two main components: an encoder network and a decoder network. As depicted in Figure 17, the encoder network takes the input data and maps it to the latent space. However, for the VAE, the final layers of the encoder produce the mean and variance parameters that describe the probability distribution of the latent space. On the other hand, the decoder network takes the sampled latent vectors and tries to reconstruct the original input data. The decoder progressively up-samples the latent vectors to match the size and complexity of the input data, ultimately producing a reconstructed output.

**Figure 17:**
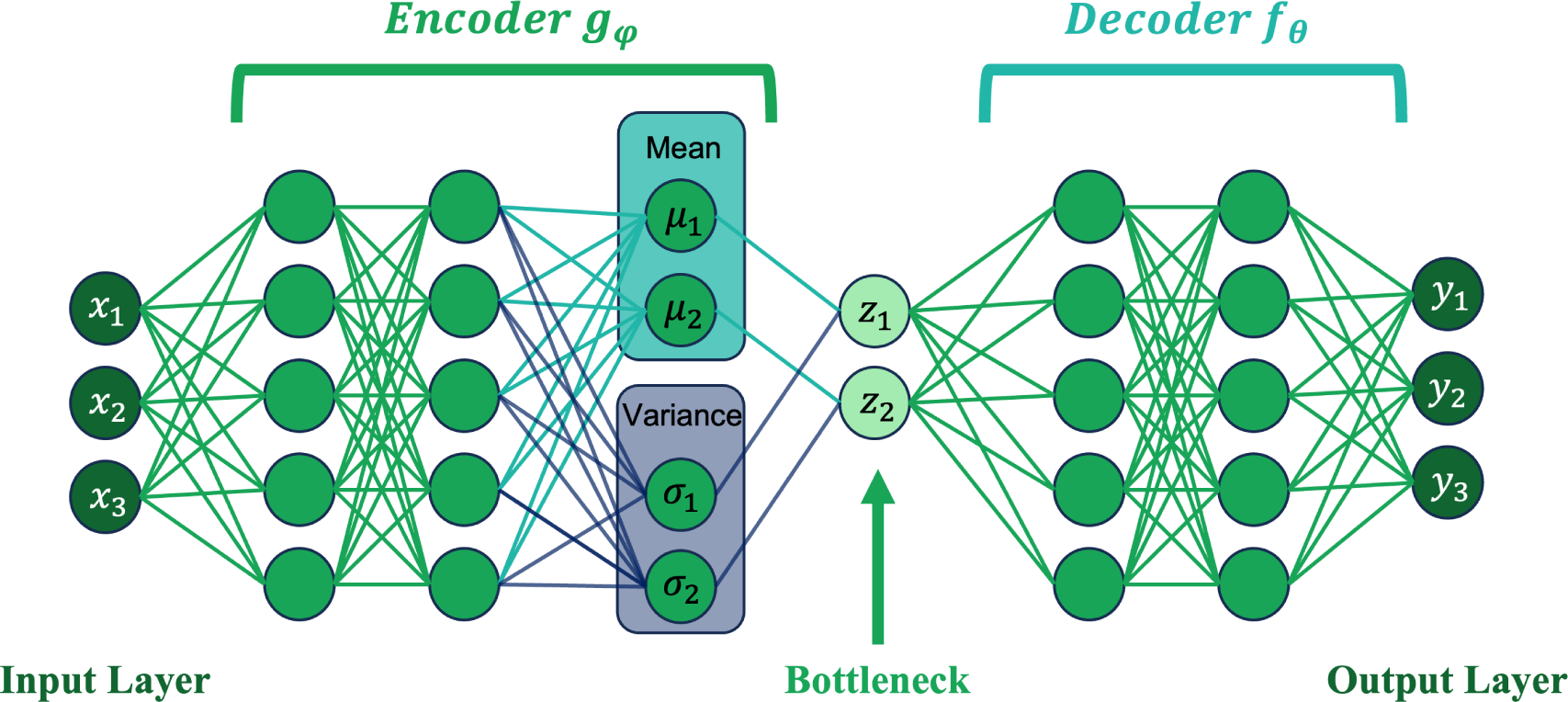
General structure of a typical Variational AutoEncoder model (VAE)

More specifically, in our study, we will reproduce and use the architecture developed in a previous study (Davidsen et al., 2019). This architecture has the advantage of having already been optimized for a similar task. In particular, it has been shown that this specific architecture is able to accurately learn some important features of the data. This architecture is illustrated in Figure 18 and consists of:

1. 3 Input layers for the CDR3 sequence, the v-gene and the j-gene
2. An embedding of each input element
3. A concatenation of the 3 elements’ embeddings into one matrix
4. Dense layers that deduce important information from the data
5. The construction of the mean and log variance of the latent space probability distribution
6. Sampling of the latent space vector
7. Dense layers of the decoder network that try to retrieve the previous inputs
8. Dissociation and decoding of each element
9. 3 Output layers for the CDR3 sequence, the v-gene and the j-gene

**Figure 18:**
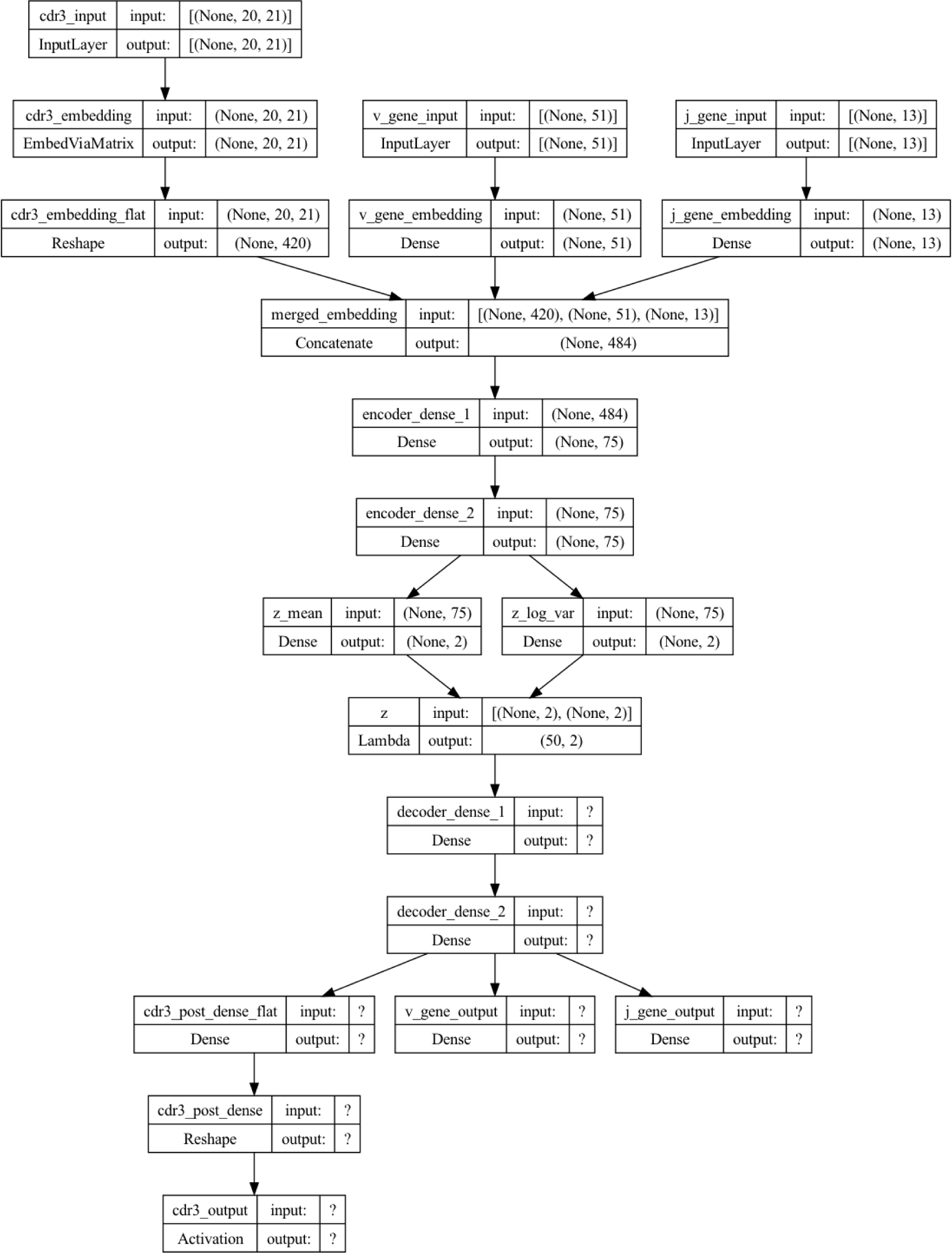
Architecture of the VAE (Davidsen et al., 2019)

#### 4.4.1 Training

The training of the VAE involves optimizing the parameters of both the encoder and decoder networks. We optimize those parameters in terms of a weighted combination of 3 objective functions for the 3 inputs and outputs of our VAE model:

- The main objective function for the CDR3 sequence *L*(*θ, ϕ*)
- The cross-entropy loss function for the v-gene 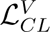
- The cross-entropy loss function for the j-gene 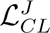

The main objective function for the CDR3 sequence is a combination of two terms: the reconstruction loss, which measures the discrepancy between the input data and the generated output, and the KL divergence, which regularizes the latent space to follow a desired prior standard Gaussian distribution. The KL-divergence between two probability distributions simply measures how much they diverge from each other. We define the objective function mathematically as follows:

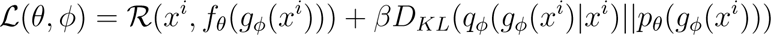

where *β* is chosen constant, *R*(*x^i^, f_θ_*(*g_ϕ_*(*x^i^*))) is the reconstruction loss and *D_KL_* is the KL-divergence:

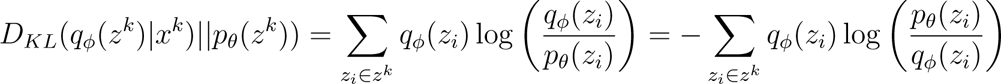

which can be simplified for the Variational AutoEncoder as it uses the Gaussian distribution (Kingma and Welling, 2022):

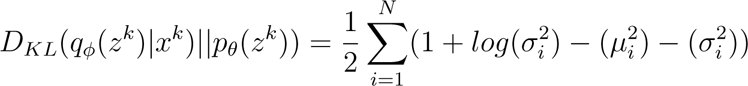

On the other hand, for this VAE model, we choose the reconstruction loss to be the cross-entropy similarly as for the v-gene and j-gene:

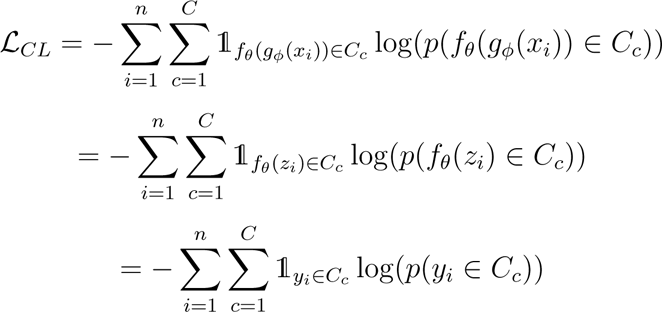

These loss functions are combined to construct a unique weighted objective function:

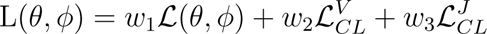

The weights used are chosen to be the optimized ones found in a previous study (Davidsen et al., 2019):

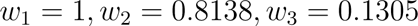

We jointly optimize our parameters *θ* and *ϕ* to minimize this objective function using as before the Adam optimizer for 100 epochs. As we can observe in Figure 19, the three losses seem to decrease likewise through the epochs. We also note that through the 100 epochs, the combined objective function for the validation set decreases significantly, thus justifying the choice of such a high number of epochs for the training of the model.

**Figure 19:**
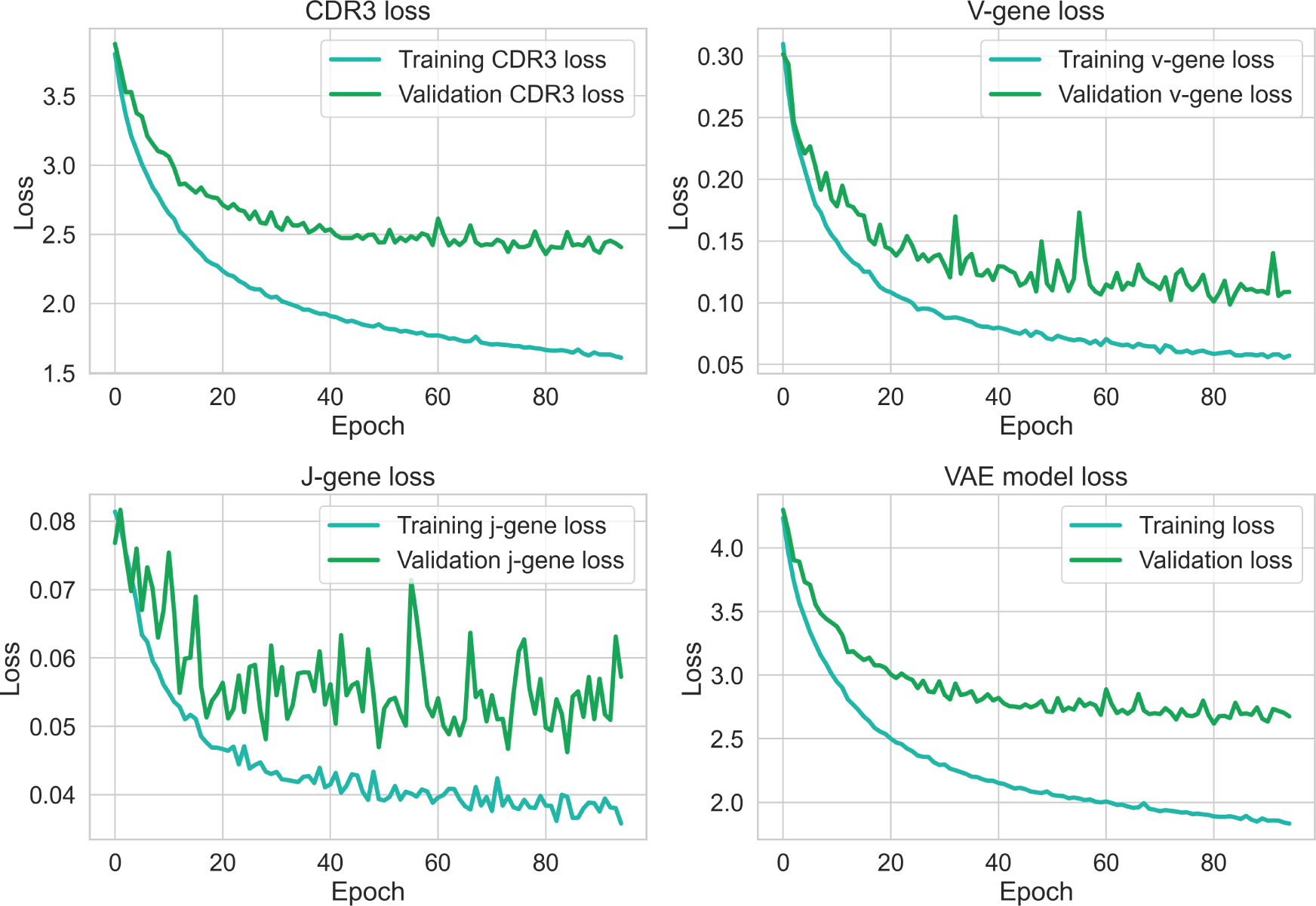
Training and validation losses of the Variational AutoEncoder model.

An critical aspect during the model training process is the computation of the relationship between each parameter in the network and the final output loss. In general, for neural networks, this calculation is achieved by a technique called back-propagation. However, applying backpropagation to a random sampling process is not feasible. Fortunately, there is a clever approach called the “reparameterization trick” that provides a solution. This trick, described in Figure 20 involves randomly sampling from a unit Gaussian distribution and then adjusting the sampled values by the mean and variance of the latent distribution. By using this reparameterisation, we can optimize the parameters of the distribution while retaining the ability to generate random samples from it.

**Figure 20:**
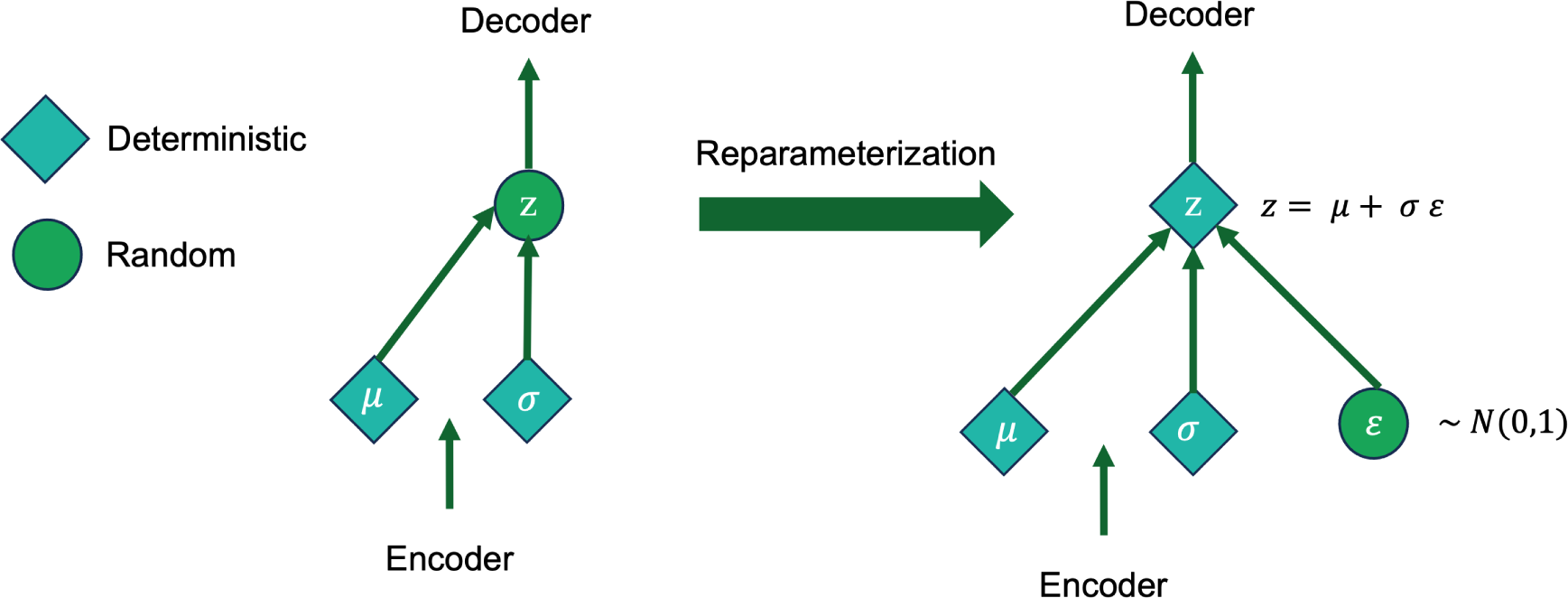
Schematization of the reparameterization trick for the VAE model.

Another technique used in our model addresses the challenge of the network learning negative values for the standard deviation value. To overcome this problem, we train the network to learn the logarithm of the standard deviation (log(*σ*)) instead.

#### 4.4.2 Choosing the entanglement parameter

In the previous section, we have defined the objective function for the VAE model using a constant *β*. This constant *β* is essential for our VAE model and is often called the *entanglement*. By varying its value between 0 and 1, we attach more or less importance to the cross-entropy loss or the KL-divergence. We have for example as in Figure 21:

**Figure 21:**
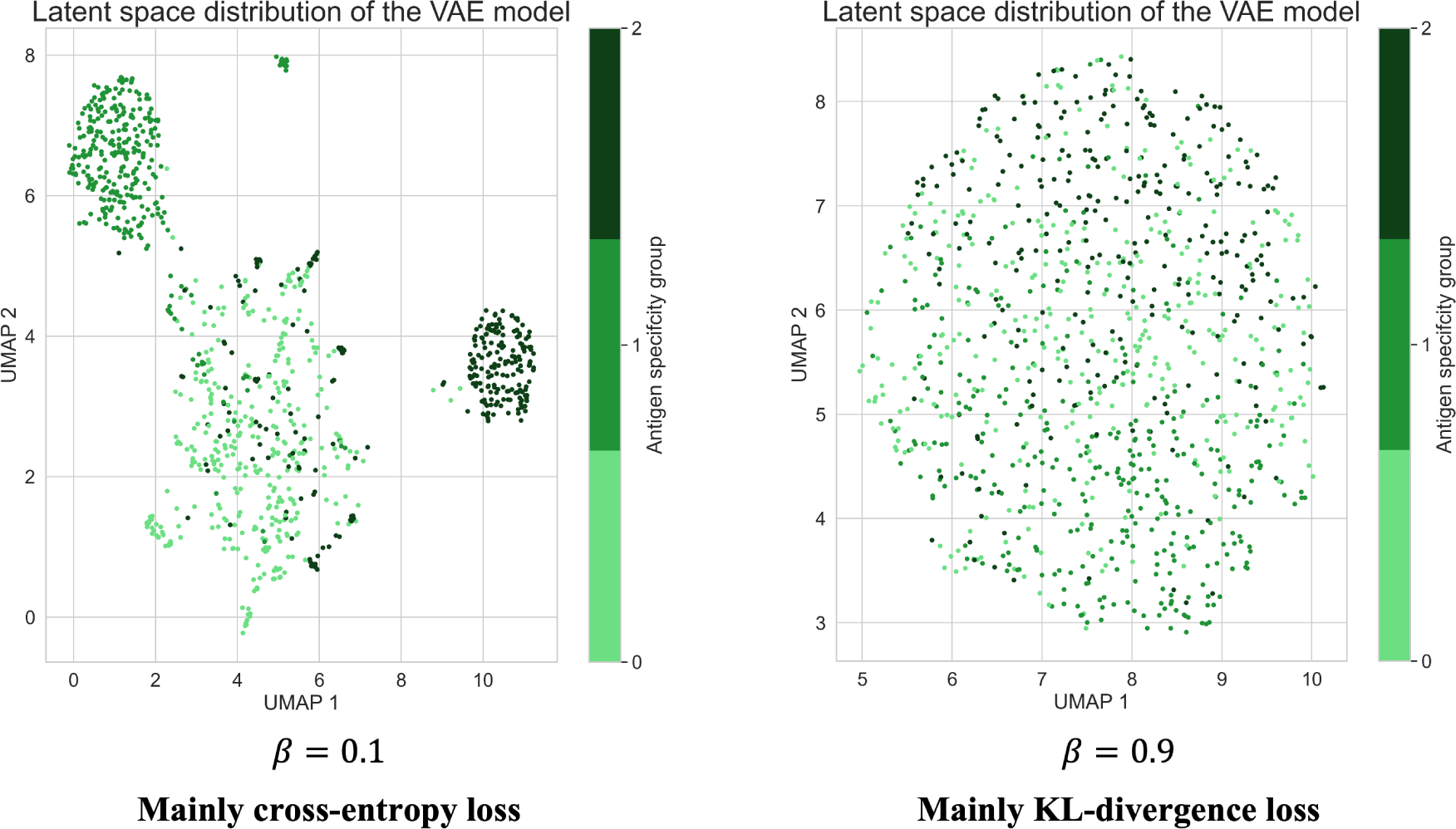
Latent space distributions for a VAE model purely optimized for recon-struction loss or for KL-divergence loss.

- If *β* is close to 0, our VAE model is mainly optimized to minimize the cross-entropy loss. Therefore, the embeddings on a 2D latent space reveal the formation of distinct clusters. However, we can observe that the latent space is highly discontinuous greatly limiting the relevancy and performance of our model.
- If *β* is close to 1, our VAE model optimizes the KL-divergence loss. Intuitively, this loss encourages the encoder to distribute all embeddings evenly around the center of the latent space. If it tries to “cheat” by clustering them apart into specific regions in the space (e.g., away from the origin) it will be penalized. This results in a model that has a poor clustering performance even though its probability distribution space seems to be centered and continuous.

On the other hand, optimizing the two losses together with a balanced objective function results in the generation of a latent space that maintains the notion of similarity or dissimilarity of nearby embeddings on the local scale via clustering. Moreover, in this setting, the embeddings are still very densely packed near the latent space origin as it is depicted in Figure 22. Therefore, choosing the correct value of *β* is essential to maintain these important performance properties. For our dataset, this corresponds to choosing a value of *β* around 0.2 (*β ≈* 0.2).

**Figure 22:**
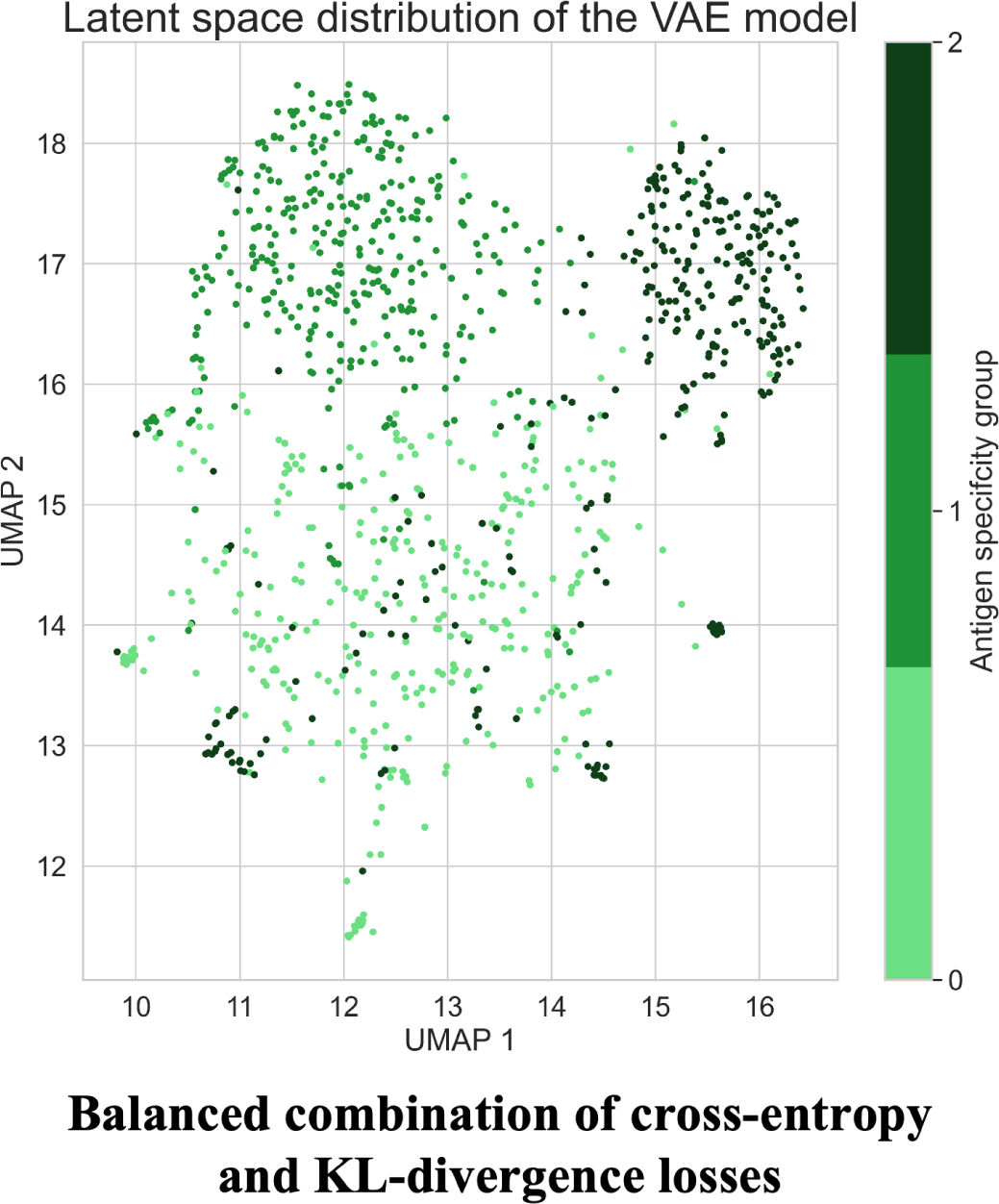
Latent space distribution for a VAE model evenly optimized for recon-struction loss and for KL-divergence loss.

### 4.5 Transfer Learning and Transformers

#### 4.5.1 Impact and benefits of Transformers and Transfer Learning

Transformer models are a type of deep learning model that has gained significant attention and popularity in Natural Language Processing (NLP) tasks. They were introduced in a 2017 paper (Vaswani et al., 2017) and have since become the backbone of many state-of-the-art NLP systems. Their ability to capture contextual relationships and dependencies within text has led to remarkable advances in tasks such as machine translation, sentiment analysis, text generation and more.

Transformers are a type of NLP models that has revolutionized the field of NLP by overcoming some of the limitations of traditional sequence-based models such as Recurrent Neural Networks (RNNs). They use a self-attention mechanism that allows them to capture dependencies between different words in a sentence or sequence. This mechanism allows the model to focus on different parts of the input sequence when making predictions, without the need for explicit sequential processing.

Like AutoEncoders or Variational AutoEncoders, transformers consist of an encoder and a decoder. The encoder takes an input sequence and encodes it into a set of hidden representations, while the decoder generates an output sequence based on these representations.

Transformers have become essential for a number of reasons:

- Parallelization: Transformers can process inputs in parallel, making them faster and more efficient, particularly for long sequences.
- Long-term Dependencies: Transformers can capture long-term dependencies. This makes them more effective in tasks that require understanding context over a span of characters.
- Attention Mechanism: The self-attention mechanism in transformers allows the model to attend to relevant parts of the input, resulting in better contextual representation and more accurate predictions.
- Transfer Learning: Transformers have played a pivotal role in transfer learning, where models pre-trained on large datasets can be fine-tuned on specific down-stream tasks with smaller datasets. This approach has significantly reduced the need for massive amounts of labeled data and improved the performance of unsupervised models.

In particular, the attention mechanism is an essential feature of the Transformers models. We can think of the attention mechanism, or self-attention, as a mechanism that enhances the information content of an input embedding by incorporating information about the input’s environment. As shown in Figure 23, this mechanism is used multiple times through *”Multi-Head Attention”* steps. Self-Attention compares all input sequence members with each other and modifies the corresponding output sequence positions. In other words, the self-attention layer differentiatively key-value searches the input sequence for each input, and adds the results to the output sequence. Mathematically, self-attention is calculated using dot products between query, key and value vectors, which are linear projections of the input sequence:

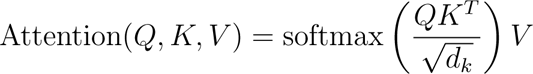

where:

1. Q, K, V are matrices packing together sets of queries, keys, and values, respectively.
2. *k* which takes the value of each of the elements of the input, corresponds to a key.
3. *d_k_* is the dimension for each key *k*.

**Figure 23:**
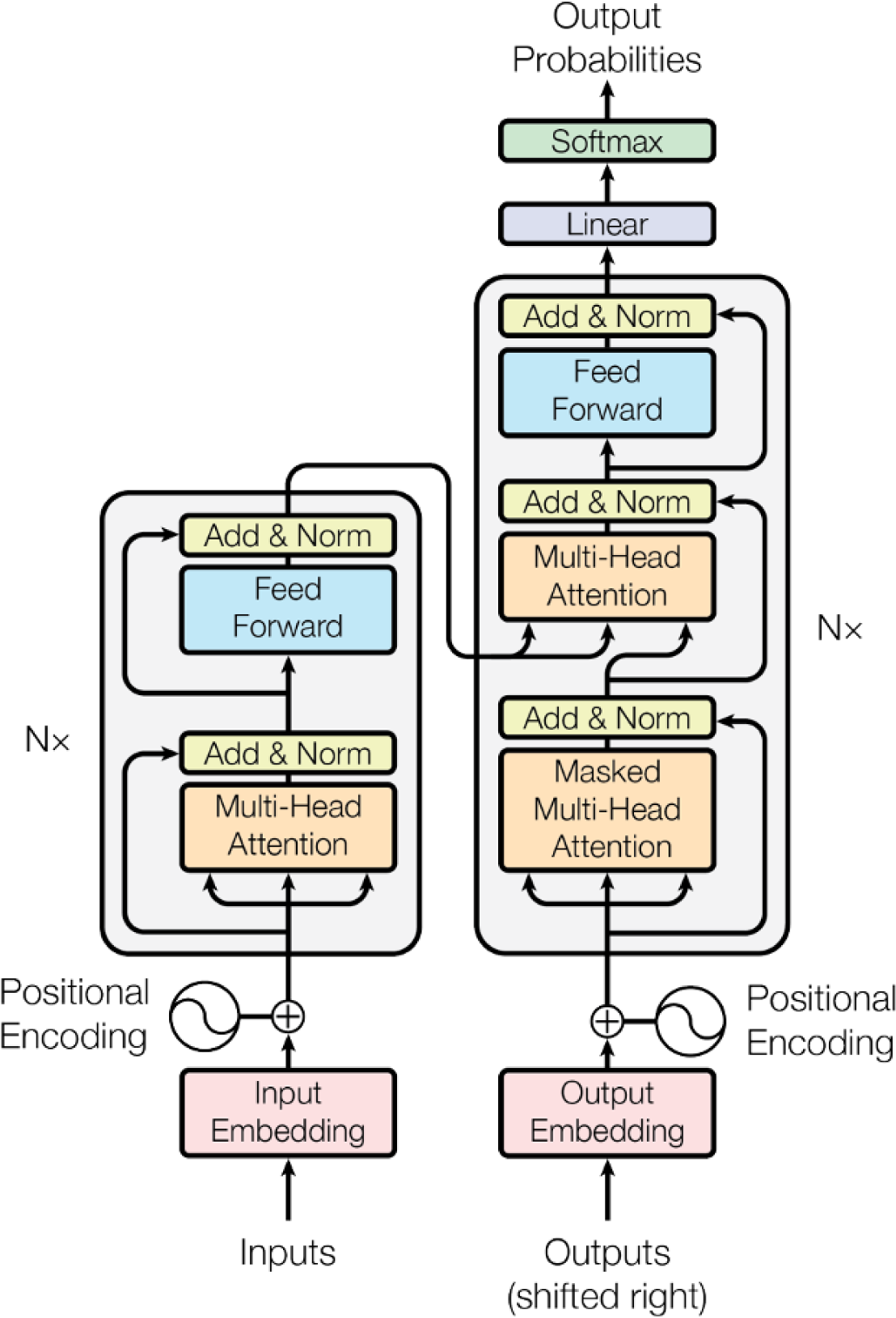
General architecture of a Transformers model (Vaswani et al., 2017)

The weighted values are then combined to form the output representation. Self-attention allows transformers to capture dependencies between different positions in the sequence, providing a flexible and powerful way to model relationships.

The multi-head attention mechanism in transformers and described in Figure 24 involves multiple linear projections of queries, keys and values. The resulting projections are processed in parallel by a single attention mechanism. These intermediate results are concatenated and projected again to produce a final result. This multi-head attention approach enhances the model’s ability to capture diverse relationships and dependencies within the input sequence, leading to more comprehensive and ex-pressive representations.

**Figure 24:**
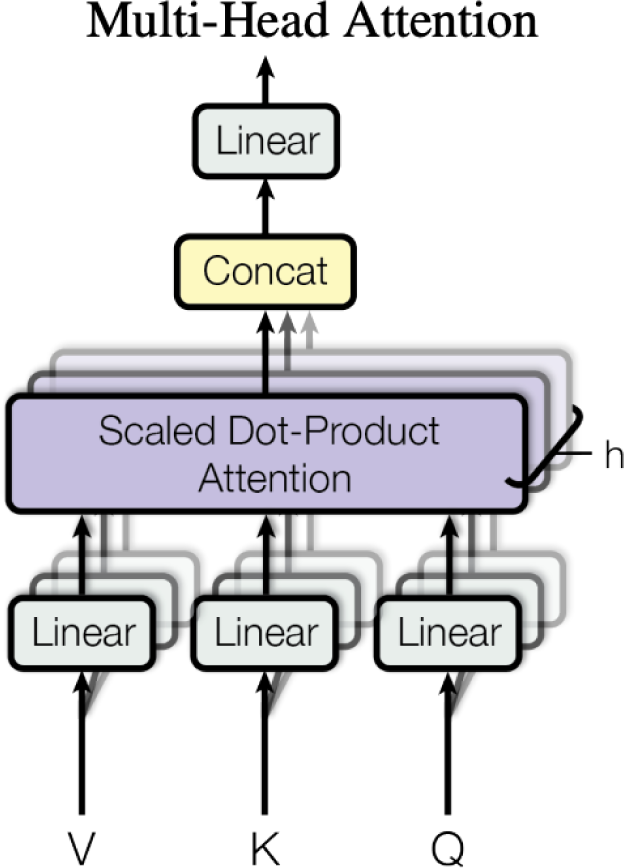
Multi-Head attention mechanism (Vaswani et al., 2017)

On the other hand, the use of transfer learning in our task can significantly improve accuracy and efficiency by leveraging knowledge from pre-trained models, reducing data requirements, and capturing complex relationships through domain-specific knowledge transfer. This approach improves the generalization, scalability and biological relevance of TCR clustering, leading to a deeper understanding of TCR repertoires and facilitating the development of personalized immunotherapies.

#### 4.5.2 TCR-BERT

Therefore, in this section we will present and use a Transformer model that mimics the usual architecture of an NLP model, but has been adapted for TCR specificity analysis: TCR-BERT (Wu et al., 2021). In fact, similar to the AutoEncoder and the Variational AutoEncoder, TCR-BERT uses unlabelled TCR sequences to learn a versatile representation of TCR sequences in a latent space. BERT (**B**idirectional **E**ncoder **R**epresentations from **T**ransformers) models excel in capturing the contextual relationships and semantic understanding of language. In the case of TCR-BERT, the pre-training process allows it to learn the grammatical structure and patterns within TCR sequences. While this does not directly encode TCR specificity information, it enables TCR-BERT to capture relevant sequence features and dependencies that may contribute to specificity. Using a similar method as before, we can then leverage UMAP to perform dimensionality reduction and K-means for clustering.

Specifically, the goal of the TCR-BERT model is to create a continuous embedding of T-cell receptor sequences that can be used for a variety of subsequent tasks. Prior to training, the TCR-BERT model learns to predict hidden amino acids based on the context around them, capturing the structural patterns in TCR sequences. Pre-training is performed on a large corpus of sequences, without regard to the understanding of antigen binding affinities. The model is then further trained to predict the antigen from a collection of 45 antigen labels to which a given amino acid sequence will bind. It is shown in this research that TCR-BERT can be used for various tasks such as antigen binding prediction and TCR clustering for in-depth TCR analysis.

We will make use of the pre-trained version **2)** in Figure 25 of TCR-BERT freely available online on the *Hugging Face* platform. Through the Hugging Face library and platform, we can easily leverage pre-trained models for various tasks. Moreover, *Hugging Face*’s user-friendly interfaces, documentation, and model repositories provide a seamless experience, enabling us to harness the power of advanced models with ease. This pre-trained version has the advantage to have already been trained on a labeled dataset.

**Figure 25:**
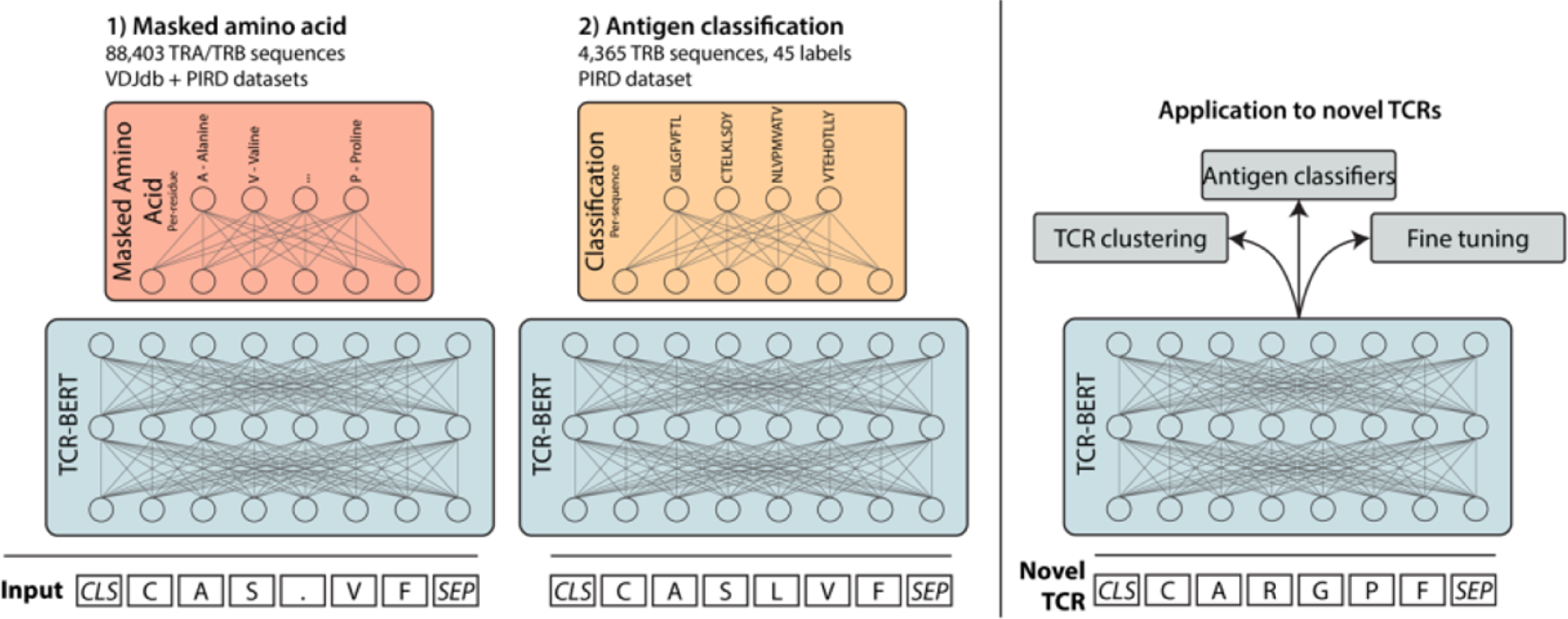
TCR-BERT self-supervised training (Wu et al., 2021)

#### 4.5.3 Pre-trained vs Fine-tuned

We now want to compare the performance of two Transformers models in the context of TCR analysis, focusing specifically on the analysis of TCR sequences associated with Sars-Cov-2. The first model we can evaluate is the pre-trained TCR-BERT without fine-tuning to assess its performance on our Sars-Cov-2 dataset. This helps us to understand how well the pre-trained TCR-BERT captures the relevant features and patterns in the TCR sequences related to Sars-Cov-2. From Figure 26, this model performs relatively well for clustering our TCR sequences. We obtain quite clear and distinct clusters with a relatively high silhouette score. In addition, we can inspect the logo plots of each cluster as they reveal the patterns in the sequences of each cluster. We notice clear patterns: sequences that end with ’EAFF’ tend to have small *UMAP 1* embeddings while sequences ending with ’EQYF’ tend to have larger *UMAP 1* embeddings. However, as the model was trained on a different dataset and is not optimized to understand well the specific structures and similarities of TCR sequences related to Sars-Cov-2, we could improve its performance further.

**Figure 26:**
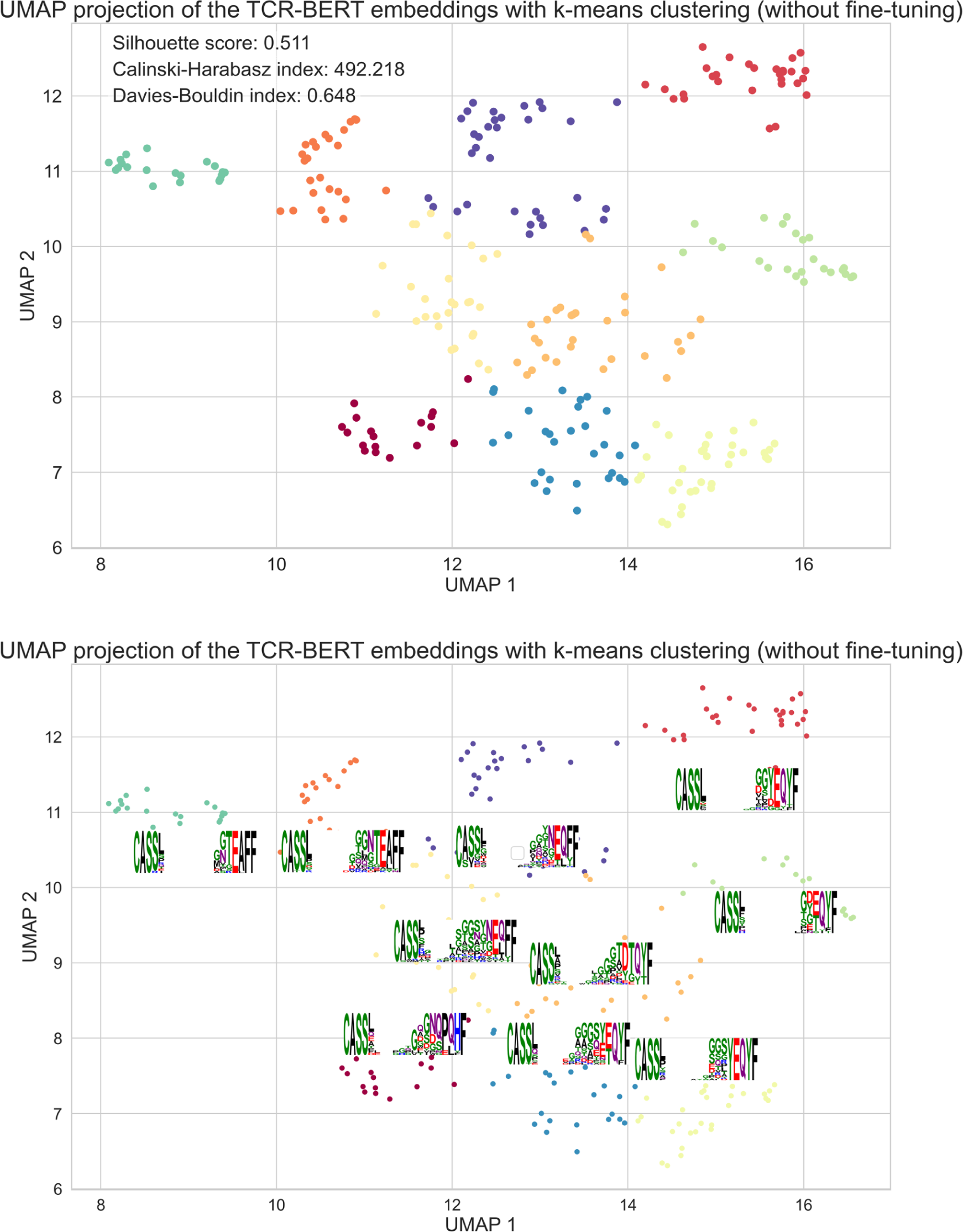
K-means clustering of the embeddings from TCR-BERT along with the corresponding logos for each cluster (without fine-tuning)

The second model we examine is the fine-tuned version of TCR-BERT using 100, 000 TCR sequences from our dataset. Indeed, fine-tuning involves training further the pre-trained TCR-BERT model using our specific Sars-Cov-2 dataset. By fine-tuning the model, we aim to enhance its performance and adapt it to the specific characteristics and nuances of the TCR sequences associated with Sars-Cov-2 while still leveraging some essential prior knowledge. We produce the same method as previously to build Figure 27. By computing some clustering performance metrics, we can deduce that this fine-tuned model would perform slightly better for our task. Moreover, we notice similar behaviors and patterns as for the pre-trained only model: sequences that end with ’EAFF’ tend to have small *UMAP 2* embeddings while sequences ending with ’EQYF’ tend to have larger *UMAP 2* embeddings.

**Figure 27:**
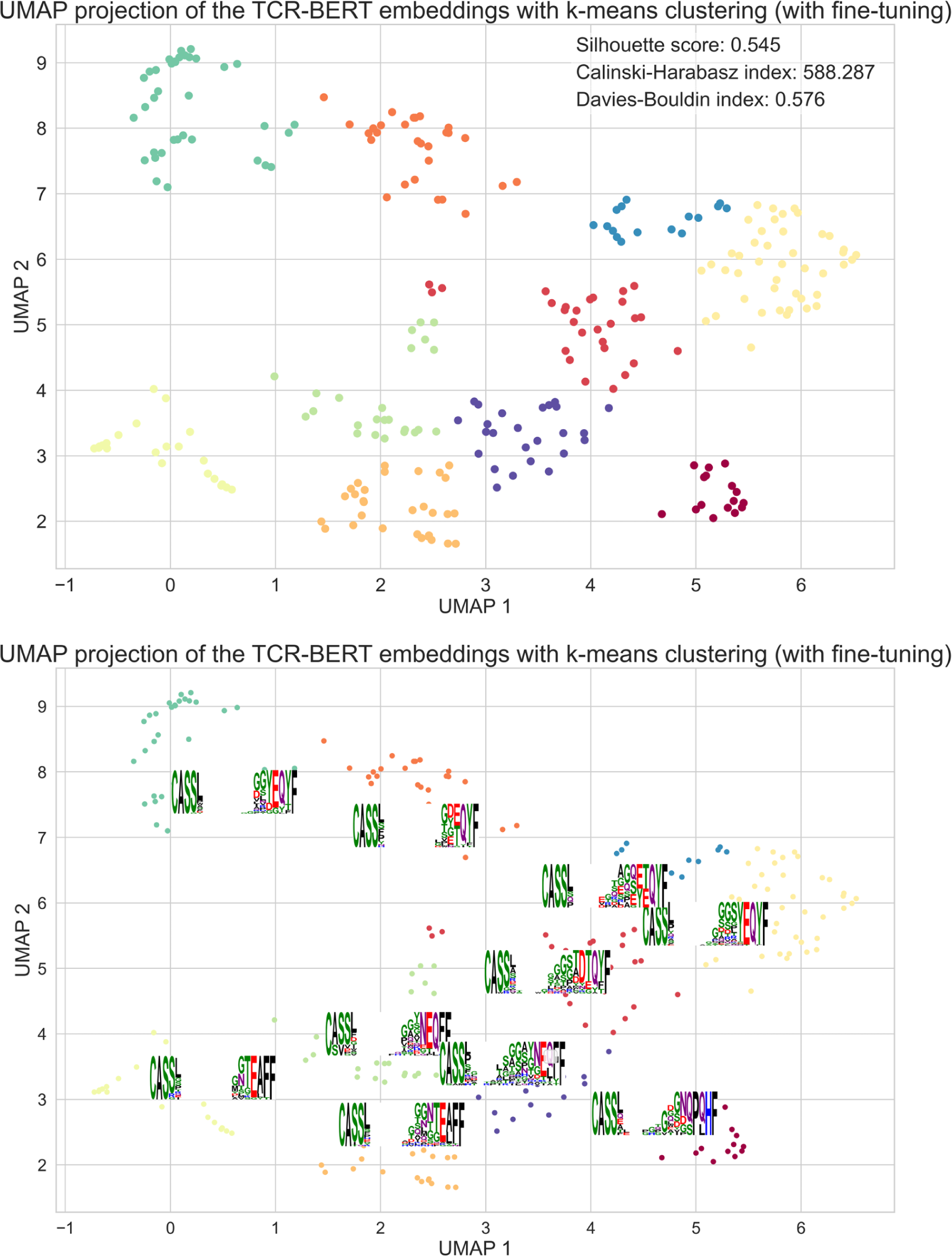
K-means clustering of the embeddings from the fine-tuned TCR-BERT along with the corresponding logos for each cluster.

By comparing the performance of the pre-trained TCR-BERT and the fine-tuned TCR-BERT model on our Sars-Cov-2 dataset, we have gained insights into the effectiveness of fine-tuning for TCR analysis tasks. Even though, the performance improvement is only small, this could have important impacts and consequences on future downstream tasks.

## Acknowledgements

I would like to express my deep gratitude and appreciation to *Dr Barbara Bravi*, for her invaluable guidance, support, and mentorship throughout the course of this research. Her expertise, dedication, and insightful feedback have been instrumental in shaping this paper and my overall growth as a researcher. I am truly grateful for her unwavering belief in my abilities and the opportunities she has given me.

Furthermore, I would like to extend my sincere thanks to the *AI for Health Lab* at Stanford University for their assistance to this study. The resources, expertise, and collaborative environment provided by *Kevin Wu* and *Kyle Swanson* have been instrumental in shaping some important parts of this research. The cutting-edge technologies and interdisciplinary discussions with the lab have enriched my understanding of the existing methods.

## A Appendix

You can find freely access here the code that has been done during the elaboration of this study.

Most of the code has been cleaned and annotated with comments explaining the methods and techniques used. In fact, reproducible code is essential for conducting rigorous and reliable research. The data analyses and scientific claims can be verified and replicated by others using the same data and software.

Please read the README.md file for more information on how to load the datasets and run the code successfully. A requirements.txt file has also been added to the GitHub repository in order to simplify the installation of all the necessary packages and avoid any conflicts.

## B Appendix

For an easier and faster training and testing of the Transformers model, here is a link to a shared Google Colab document. It already contains the necessary code to run the training and fine-tuning of the model using a custom dataset. This implementation allows anyone to make use of GPU computational resources for an accelerated training. Please note that you have to select *GPU runtime* in the *Runtime* parameters of the notebook.

## C Appendix

Moreover, you can also find here a web application simulating some TCR-antigen sequences binding examples and their predictions based on the chosen model (AutoEn-coder, Variational AutoEncoder, etc…): https://m4r-dash.yanismiraoui.repl.co/. You can also ask any questions that you may have about this research paper to a Chat-bot. The code of this web application is available here and has been deployed and hosted on a standard virtual machine using Heroku, Replit and Amazon Web Ser-vices (AWS).

Please note that as this web application is hosted with limited resources, some results and figures may take some time to load.

On the other hand, the Figures 14 and 16 were created using this diagram creation website. It greatly facilitated the drawing of the full neural networks architectures along with the sizes of each layer.

## Footnotes

1 In this context, the term “*automatic*” refers to a method or approach that uses computational tools and algorithms to identify the specific TCRs involved in the immune response to COVID-19.

3 lease note that we have selected only the columns meaningful for our analysis for this visu-alisation. The original datasets contains 52 columns

4 Groups of TCRs that are biochemically similar

5 GitHub repository: https://github.com/statbiophys/OLGA

